# The circulating SARS-CoV-2 spike variant N439K maintains fitness while evading antibody-mediated immunity

**DOI:** 10.1101/2020.11.04.355842

**Authors:** Emma C. Thomson, Laura E. Rosen, James G. Shepherd, Roberto Spreafico, Ana da Silva Filipe, Jason A. Wojcechowskyj, Chris Davis, Luca Piccoli, David J. Pascall, Josh Dillen, Spyros Lytras, Nadine Czudnochowski, Rajiv Shah, Marcel Meury, Natasha Jesudason, Anna De Marco, Kathy Li, Jessica Bassi, Aine O’Toole, Dora Pinto, Rachel M. Colquhoun, Katja Culap, Ben Jackson, Fabrizia Zatta, Andrew Rambaut, Stefano Jaconi, Vattipally B. Sreenu, Jay Nix, Ruth F. Jarrett, Martina Beltramello, Kyriaki Nomikou, Matteo Pizzuto, Lily Tong, Elisabetta Cameroni, Natasha Johnson, Arthur Wickenhagen, Alessandro Ceschi, Daniel Mair, Paolo Ferrari, Katherine Smollett, Federica Sallusto, Stephen Carmichael, Christian Garzoni, Jenna Nichols, Massimo Galli, Joseph Hughes, Agostino Riva, Antonia Ho, Malcolm G. Semple, Peter J.M. Openshaw, J. Kenneth Baillie, The ISARIC4C Investigators, the COVID-19 Genomics UK (COG-UK) consortium, Suzannah J. Rihn, Samantha J. Lycett, Herbert W. Virgin, Amalio Telenti, Davide Corti, David L. Robertson, Gyorgy Snell

## Abstract

SARS-CoV-2 can mutate to evade immunity, with consequences for the efficacy of emerging vaccines and antibody therapeutics. Herein we demonstrate that the immunodominant SARS-CoV-2 spike (S) receptor binding motif (RBM) is the most divergent region of S, and provide epidemiological, clinical, and molecular characterization of a prevalent RBM variant, N439K. We demonstrate that N439K S protein has enhanced binding affinity to the hACE2 receptor, and that N439K virus has similar clinical outcomes and *in vitro* replication fitness as compared to wild-type. We observed that the N439K mutation resulted in immune escape from a panel of neutralizing monoclonal antibodies, including one in clinical trials, as well as from polyclonal sera from a sizeable fraction of persons recovered from infection. Immune evasion mutations that maintain virulence and fitness such as N439K can emerge within SARS-CoV-2 S, highlighting the need for ongoing molecular surveillance to guide development and usage of vaccines and therapeutics.

## INTRODUCTION

SARS-CoV-2, the cause of COVID-19, emerged in late 2019 and expanded globally, resulting in over 41 million confirmed cases as of October 2020. Molecular epidemiology studies across the world have generated over 135,000 viral genomic sequences and have been shared with unprecedented speed via the GISAID Initiative (https://www.gisaid.org/). These data are essential for monitoring virus spread (Meredith et al., 2020) and evolution. Of particular interest is the evolution of the SARS-CoV-2 surface protein, spike (S), which is responsible for viral entry via its interaction with the human angiotensin-converting enzyme 2 (hACE2) receptor on host cells. The S protein is the target of neutralizing antibodies generated by infection (Jiang et al., 2020) or vaccination (Folegatti et al., 2020; Jackson et al., 2020; Keech et al., 2020) as well as monoclonal antibody (mAb) drugs currently in clinical trials (Hansen et al., 2020; Jones et al., 2020; Pinto et al., 2020).

A SARS-CoV-2 S variant, D614G, is now dominant in most places around the globe (Callaway, 2020). Studies *in vitro* indicate that this variant may have greater infectivity while molecular epidemiology indicates that it spreads efficiently and likely maintains virulence (Hu et al., 2020; Korber et al., 2020; Volz et al., 2020; Zhang et al., 2020). Amino acid 614 is outside the receptor binding domain (RBD) of S, the domain targeted by 90% of neutralizing antibody activity in serum of SARS-CoV-2 survivors (Piccoli et al., 2020). Initial studies suggest that D614G actually exhibits increased sensitivity to neutralizing antibodies, likely due to its effects on the molecular dynamics of the spike protein (Hou et al., 2020; Yurkovetskiy et al., 2020). Therefore, this dominant variant is unlikely to escape antibody-mediated immunity.

The low numbers of novel mutations reaching high frequency in sequenced SARS-CoV-2 isolates may relate to the moderate intrinsic error rate of the replication machinery of SARS-CoV-2 (Li et al., 2020c; Robson et al., 2020) and to this new human coronavirus requiring no significant adaption to humans (MacLean et al., 2020). Nevertheless, the increasing number of infected individuals and the large reservoir of individuals susceptible to infection increases the likelihood that novel variants that impact vaccine and therapeutic development will emerge and spread. Moreover, the full impact of immune selection, which can drive variant selection, likely has not yet had a dominant influence on the pandemic, since herd immunity has not yet been attained. As population immunity increases and vaccines are deployed at scale this might change. The potential for circulating viral variants to derail promising vaccine or antibody-based prophylactics or treatments, even in the absence of selective pressure from the drug or vaccine, is demonstrated by the failure of a Phase III clinical trial of a mAb targeting the respiratory syncytial virus (Simoes et al., 2020), and the need for new influenza vaccines on a yearly basis. It is therefore critical to understand whether and how SARS-CoV-2 may evolve to evade antibody-dependent immunity.

Here, we examined the immunodominant SARS-CoV-2 receptor binding motif (RBM), the primary target of the neutralizing Ab response within the RBD (Piccoli et al., 2020) and found it to be less conserved than the RBD or the entire spike protein in circulating viruses. To understand the implications of this structural plasticity for immune evasion, we defined the clinical and epidemiological impact, the molecular features, and the immune response to an RBM variant, N439K. This variant has arisen independently twice, in both cases forming lineages of more than 500 sequences. As of October 2020, it has been observed in 12 countries and is the second most commonly observed RBD variant worldwide. We find that the N439K mutation is associated with a similar clinical spectrum of disease and slightly higher viral loads *in vivo* compared with isolates with the wild-type N439 residue, and that it results in immune escape from polyclonal sera from a proportion of recovered individuals and a panel of neutralizing mAbs. N439K provides a sentinel example of immune escape, indicating that RBM variants must be evaluated when considering vaccines and the therapeutic or prophylactic use of mAbs. Long term control of the pandemic will require systematic monitoring of immune escape variants and selection of strategies that address the variants circulating in targeted populations.

## RESULTS

### The RBM is the least conserved region in the SARS-CoV-2 spike protein

Competing pressures influence the evolution of the spike RBM. First, the RBM mediates viral entry (Shang et al., 2020; Walls et al., 2020; Wrapp et al., 2020b) and therefore it must maintain sufficient affinity to engage the entry receptor hACE2. Second, it is a major target of neutralizing antibodies (Robbiani et al., 2020; Rogers et al., 2020; Wec et al., 2020) and could be a primary location for the emergence of immune escape mutations. We set out to understand these competing pressures by evaluating the landscape of RBM sequence divergence observed in circulating SARS-CoV-2 variants and in other viruses of the *Sarbecovirus* lineage.

We used published X-ray structures of SARS-CoV and SARS-CoV-2 RBD:hACE2 complexes (Lan et al., 2020; Li et al., 2005) to define the RBM residues using a 6 Å distance cutoff (**Figures 1A-C**). We evaluated ∼130,000 SARS-CoV-2 genomic sequences deposited in GISAID as of October 7, 2020 and observed a high number of variants occurring in the RBM (**Figure 1A**). To understand how the divergence of the RBM compares to the divergence of the entire RBD and the whole spike protein, we divided the spike protein into three non-overlapping regions: the RBM, the RBD outside of the RBM, and the full S protein outside of the RBD. We counted individual variants occurring at least ten times, and quantified substitutions of different amino acids at the same position as separate variants. We found that the RBM is the least conserved region of S (**Figure 1B**).

**Figure 1.**
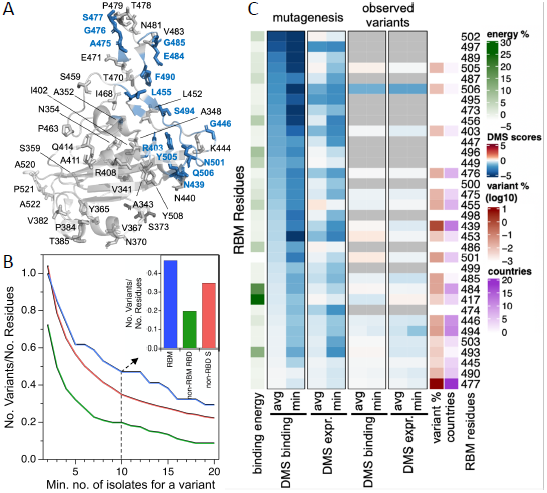
The RBM exhibits significant natural diversity in circulating SARS-CoV-2 isolates (GISAID, Oct 7, 2020, n≈130,000). **(A)** RBD variants with a minimum of 10 observed isolates in the RBM (blue) and outside the RBM (gray) mapped onto an X-ray structure of the SARS-CoV-2 RBD (PDB ID: 6M0J). **(B)** Number of observed variants in three non-overlapping S regions (RBM, non-RBM RBD, non-RBD S) normalized by the total number of residues in each region, where the number of observed isolates required to define a variant is varied. Inset: barplot representation where at least 10 isolates are observed per variant. **(C)** Heatmap of Deep Mutational Scanning (DMS) hACE2 binding and expression data for RBM residues (Starr et al., 2020). DMS score is the binding or expression fold change over WT on a Log10 scale. Aggregated DMS data is shown for each residue by taking the minimum (most disruptive variant) or the average score across all possible variants of a residue, except for the reference residue and the stop codon (‘mutagenesis’ columns). Alternatively, minimum and average scores are computed only across variants that have naturally occurred (‘observed variants’ columns). When no natural variants have been observed, cells are grey. The heatmap is annotated with frequency of non-reference amino acids in deposited sequences (at least 4 sequences were required to call a variant), in Log10 scale; number of countries in which a variant was observed; and percentage of total binding energy between RBD and hACE2 computed from an X-ray crystal structure. Data were sorted on the leftmost DMS column.

To understand this result further, we evaluated a published deep mutational scanning (DMS) data set of the RBD (Starr et al., 2020) and compared it to sequences of circulating viruses. The DMS data defines the effect of each possible single amino acid change on both expression of the RBD and its capacity to bind hACE2. For each position in the RBM, we compared the DMS results for all amino acid substitutions at that position versus only substitutions that have been observed in circulating SARS-CoV-2 isolates (**Figure 1C**). A subset of residues shows the largest loss of hACE2 binding upon mutation (top ∼1/3 of RBM residues in **Figure 1C**) and, as would be expected, few natural variants of these residues have been observed to be circulating to date. Surprisingly, these conserved residues each contribute weakly to the RBD:hACE2 total interaction energy (the sum of pair-wise interaction energies for all residues at the binding interface in the X-ray structure; “binding energy” in **Figure 1C**). For the majority of the RBM (bottom ∼2/3 of RBM residues in **Figure 1C**), variation in circulating virus sequences confirms the tolerance to mutation predicted by the DMS data. Notably, several RBM residues forming the strongest interactions with the receptor, e.g. K417 and E484, are not highly conserved despite their predicted importance. These results suggest that the RBM has a degree of structural plasticity whereby it is able to accommodate mutations without disrupting hACE2 binding.

Evolutionary analysis of *Sarbecoviruses* provides further support for RBM plasticity (Boni et al., 2020; Li et al., 2020b; Rambaut et al., 2020). The SARS-CoV RBM is highly divergent from the SARS-CoV-2 RBM (**Figure S1A-B**) while maintaining hACE2 binding affinity. Additionally, there are many sequence changes in the RBM across a panel of related coronaviruses from animal isolates (**Figure S1A-B, Table S1)**. To determine the ability of members of the *Sarbecovirus* lineage to bind hACE2, we produced nine recombinant RBD proteins corresponding to seven animal isolates, SARS-CoV-2, and SARS-CoV and evaluated their binding to recombinant hACE2 (**Figure S1C**). We found that three of the RBDs from animal isolates showed strong affinity for hACE2: GD Pangolin, which has a highly similar RBM to SARS-CoV-2, and GX Pangolin and Bat CoV WIV1, which have highly divergent RBMs (**Figure S1A-B**). This further supports the conclusion that the RBM is structurally plastic, while retaining binding with hACE2 as a receptor. Given this plasticity, we next considered whether an RBM variant can lead to immune evasion while retaining virulence.

### N439K is a prevalent SARS-CoV-2 RBM variant with increased ACE2 affinity

The two most commonly observed circulating RBD variants as of October 2020 contain mutations in the RBM (S477N and N439K). We first identified the N439K variant in March 2020, circulating in Scotland from lineage B.1 (Rambaut et al., 2020) on the background of D614G (Da Silva Filipe et al., 2020). Using phylogenetic analysis, we determined this variant represented a single lineage (**Figure 2A**) that increased in frequency to 553 sequences by June 20, 2020 (∼10% of the available Scottish viral genome sequences for this time period). Numbers of N439K and all other isolates decreased in Scotland concurrent with control of the pandemic by initiation of stringent public health measures and this lineage has not been detected in Scotland after June. However, the N439K variant has been observed in >659 sequences in a second lineage in Europe, first sampled in Romania on May 13, 2020, then Norway on June 23, 2020 and is now circulating in 12 countries, as well as arising independently in the U.S. (**Figure 2A-C**). As of Oct 6, 2020, all 1201 N439K variants arose from a C-to-A transversion in the third codon position, though these counts are heavily influenced by sampling frequency which varies widely between countries. As Scotland has a high sampling frequency for its population size (∼5.5M), it is possible to calculate a growth rate (Voltz and Frost, 2017) based on a comparison of the Scottish lineages. We find that the growth rate is similar to what has already been shown for the D614G background with no evidence for a faster rate of growth than N439 lineages (**Figure S2A**).

**Figure 2.**
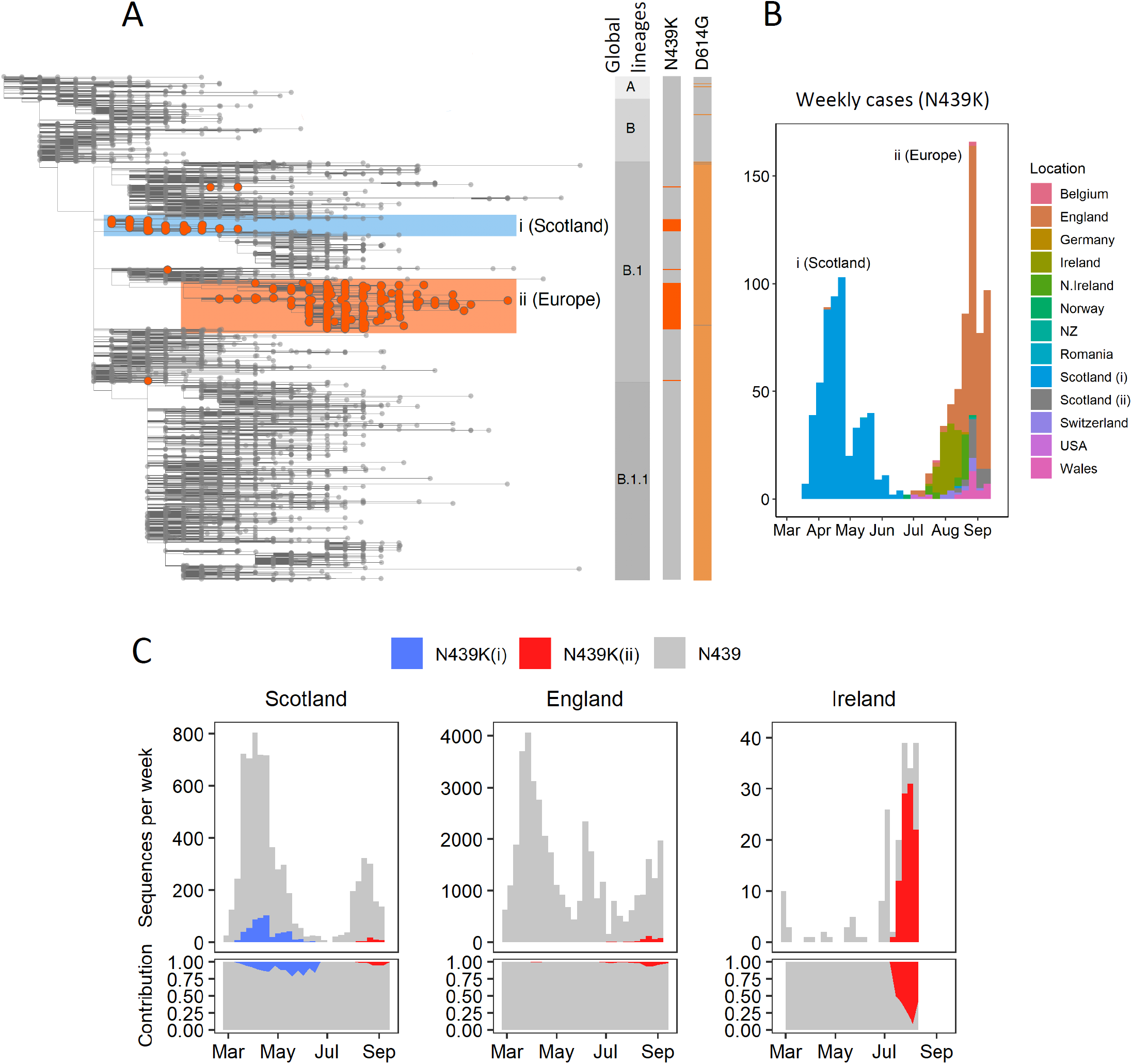
The N439K RBM variant has arisen independently multiple times, twice forming significant lineages. **(A)** Phylogenetic tree showing relationship between all global SARS-CoV-2 variants as of Oct 6, 2020, with N439K variants highlighted in orange circles. Two significant N439K lineages, one in Scotland (blue box) and one in multiple European countries (light orange box), each >500 sequences, have been detected to date. The N439K mutation has also been detected independently in the US in four linked infections and in several incidental infections. Vertical bars indicate global lineage, presence of N439K (orange), or presence of D614G (light orange). **(B)** Growth of the N439K lineages relative to sampling time and their country of occurrence. The emergence of the European lineage in Scotland is denoted as Scotland (ii). **(C)** Growth of the two N439K lineages over time (top panels) and their relative contributions (lower panels) in Scotland, England and the Republic of Ireland.

In addition to its frequency and spread, the N439K variant stood out from other circulating RBM variants as having a plausible mechanism for maintenance of viral fitness. The equivalent position to N439K in the SARS-CoV RBM is also a positively-charged amino acid (R426), which forms a salt bridge with hACE2 (Li et al., 2005). We therefore hypothesized that the N439K SARS-CoV-2 variant may form this additional salt bridge at the RBD-hACE2 interface (RBD N439K:hACE2 E329). Structural modeling supported that this salt bridge could form without disrupting the binding interface, including the two original salt bridges (RBD K417:hACE2 D30 and RBD E484:hACE2 K31) (**Figure 3A-C**). A salt bridge is the strongest type of non-covalent bond and the N439K mutation could plausibly increase the number of salt bridges at the binding interface from two to three, presenting the hypothesis that the N439K variant may have enhanced binding for hACE2.

**Figure 3.**
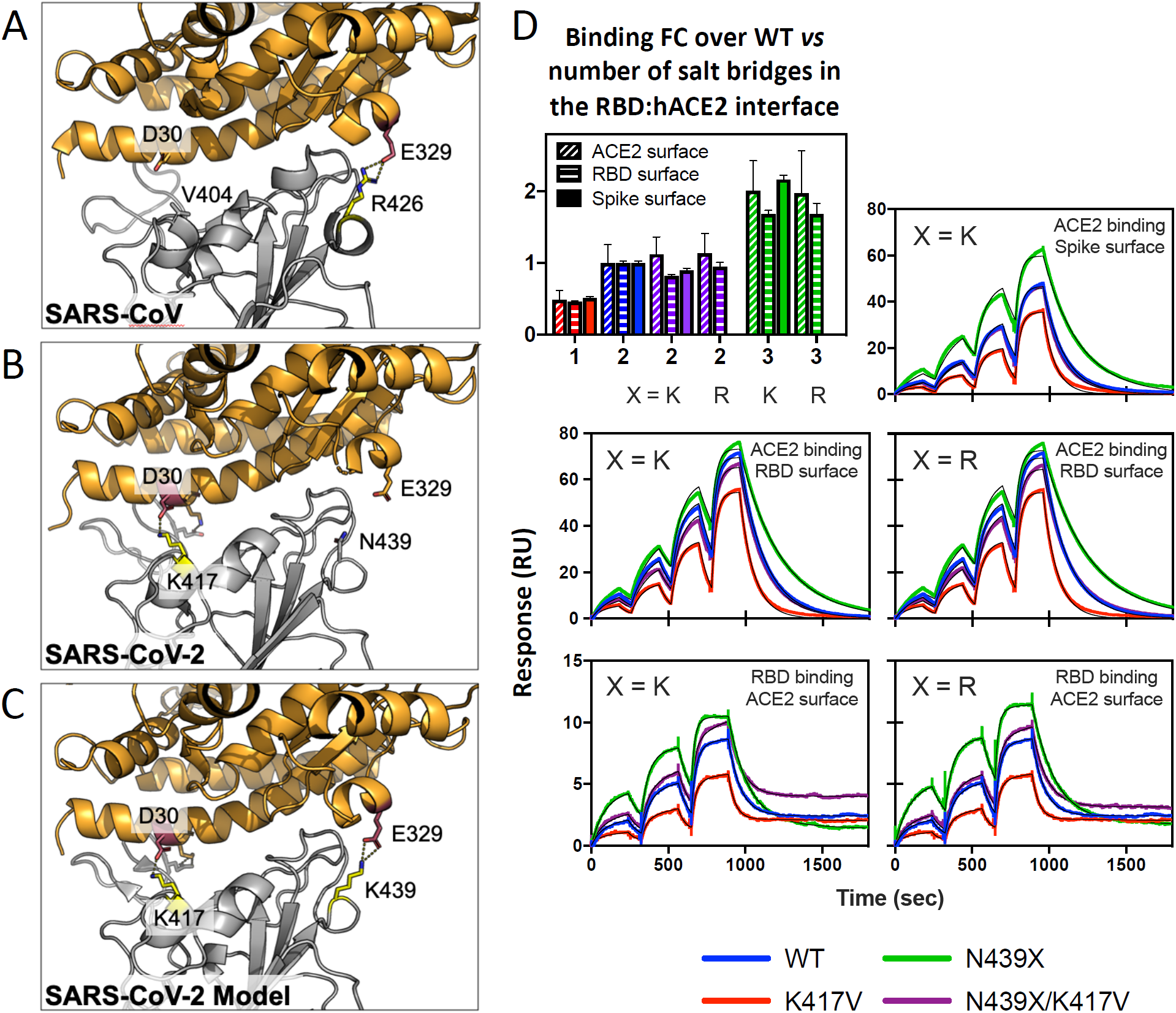
N439K enhances RBD affinity for hACE2, likely by formation of a new salt bridge at the RBD:hACE2 binding interface. **(A-B)** X-ray structures of the (A) SARS-CoV and (B) SARS-CoV-2 RBD in complex with hACE2 (based on 2AJF and 6M0J, respectively). Interface salt bridge residues are shown as sticks. SARS-CoV-2 salt bridge RBD E484:hACE2 K31 is not labeled. hACE2 is shown in orange and RBD in gray. **(C)** SARS-CoV-2 RBD in complex with hACE2 highlighting the observed K417:D30 salt bridge and the putative K439:E329 salt bridge, which was modeled in-silico. **(D)** Binding affinity of RBD and Spike variants for hACE2 measured by surface plasmon resonance. Monomeric hACE2 is injected successively at 11, 33, 100 and 300 nM onto surface-captured spike ECD or RBD; alternately, RBD is injected successively at 3.1, 12.5 and 50 nM onto surface-captured hACE2. All spike ECD contain the D614G mutation. Bar graph – Affinity measurements (averages of 3-4 replicates) expressed as a fold change relative to WT binding within each experiment format. WT K_D_ values measured as: 95 ± 1.6 nM (Spike surface), 63 ± 1.0 nM (RBD surface), 19 ± 3.3 nM (ACE2 surface); errors are SEM.

To test this hypothesis, we used surface plasmon resonance (SPR) to evaluate binding of recombinant N439K S or RBD protein to recombinant hACE2. We also evaluated the N439R and K417V variants, each of which is found in SARS-CoV at these positions. Across multiple assay formats, we found that the N439K and N439R variants exhibited a ∼2-fold enhanced binding affinity for hACE2 as compared to the original N439 variant (termed herein WT) (**Figure 3D**). The magnitude of this enhancement was paralleled by a ∼2-fold loss of binding affinity for the K417V variant relative to WT. Lastly, we also tested the effect of the N439K/R and K417V mutations in combination. These double variants form the same number of salt bridges at the hACE2 binding interface as compared to WT, but one is at RBD position 439 rather than 417; we found they had an hACE2 affinity similar to the WT (**Figure 3D**). These data indicate that acquisition of the N439K mutation enhances binding affinity, which could have implications *in vivo* in the context of natural infection. Also, the enhanced affinity could plausibly compensate for other mutations that would otherwise be detrimental (e.g. K417V), further highlighting the plasticity of the RBM.

### N439K SARS-CoV-2 maintains fitness and virulence

The enhanced hACE2 affinity of the N439K variant, its geographical emergence as independent lineages as well as its prevalence among circulating viral isolates is consistent with maintained viral fitness. We set out to directly examine fitness by evaluating clinical data and outcomes of virus carrying the N439K mutation versus WT N439, as well as by direct *in vitro* viral growth and competition.

We used qPCR to evaluate viral load (as measured by cycle threshold, Ct) in 1,918 Scottish patients whose viral isolates had been sequenced (**Figures 4A-B**). Viral isolates were either N439K/D614G (n=406), N439/D614G (n=978) or ancestral (N439/D614) (n=534). Our analysis found strong evidence that the N439K/D614G genotype was associated with marginally lower cycle threshold (Ct) than the N439/D614G genotype (mean Ct value difference between N439K/D614G and N439/D614G: -0.65, 95% CI: -1.22, -0.07) (**Figure 4B**). As Ct measurements were carried out in multiple sites, a sub-analysis of viral load using RNA standards was carried out with available samples and showed a near-complete correlation with Ct (**Figure 4B**). D614G has previously been associated with higher viral loads/lower Ct values than D614 (Korber et al., 2020) but we did not detect this difference in this statistical analysis due to the intercept of the model being imprecisely estimated (**Table S2**).

**Figure 4.**
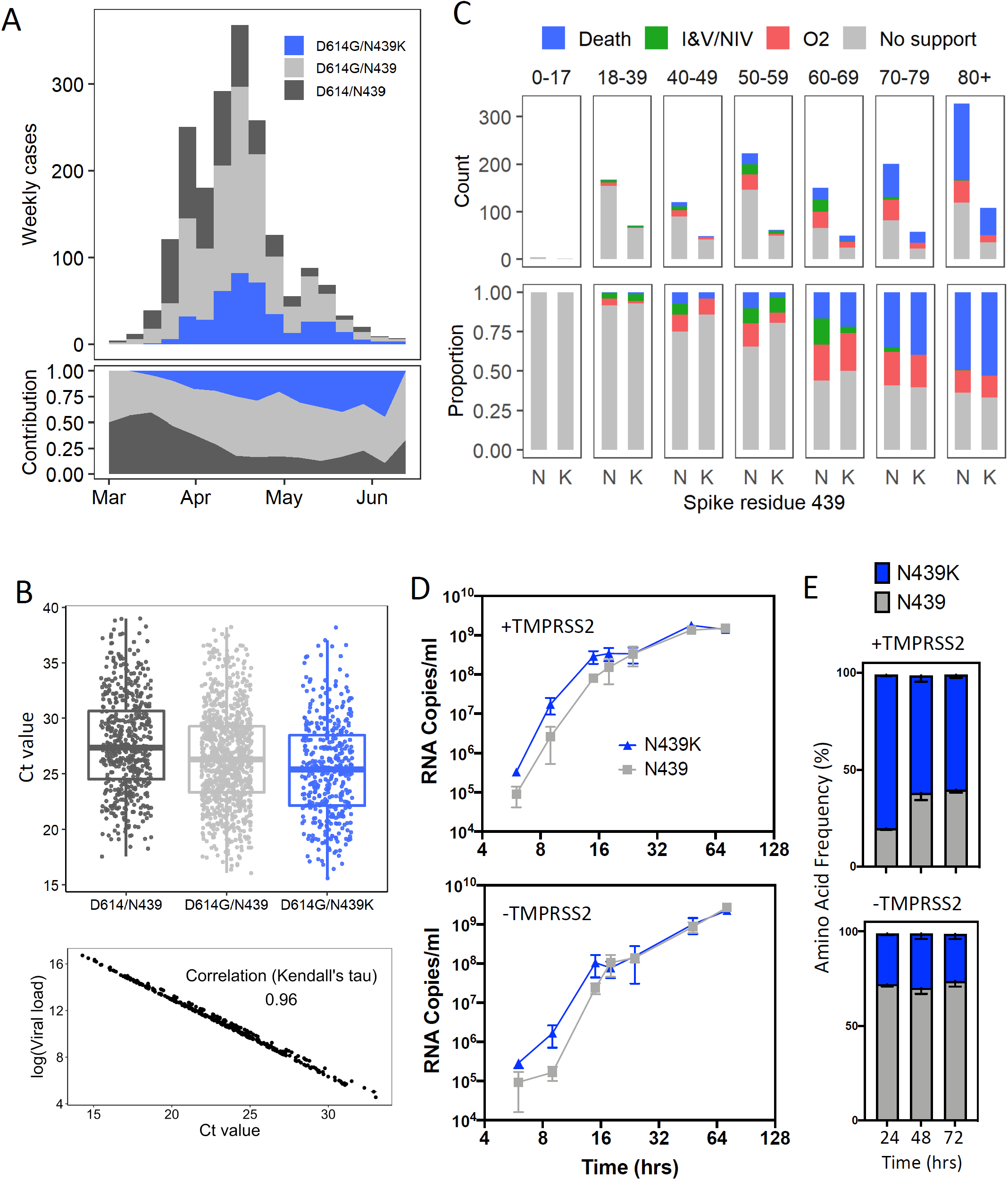
Clinical outcomes and virological evaluation of the ‘Scottish’ N439K lineage indicate maintenance of fitness relative to WT virus. **(A)** Epidemiological growth of the D614/N439, D614G/N439 or D614G/N439K virus in the NHS Greater Glasgow and Clyde Health Board area (NHS GGC) relative to sampling time in epi weeks (top) and their relative contributions (bottom) for 1918 patients whose diagnostic samples were sequenced. **(B)** Top – Real-time PCR data for N439/D614, N439/D614G and N439K/D614G groups, same patient population as in (A). The N439K genotype was associated with marginally lower Ct values than the N439 genotype (posterior mean Ct value difference between N439K/D614G and N439/D614G: -0.65, 95% CI: -1.22, -0.07). Bottom – Correlation between Ct and quantitative viral load. **(C)** Severity of disease within NHS GGC for a subset of 1591 patients. Ordinal scale scored by requirement for supplementary oxygen: 1. No respiratory support, 2: Supplemental oxygen, 3: Invasive or non-invasive ventilation or oxygen delivered by high-flow nasal cannulae, 4: Death. Analysis based on the ordinal scale indicated that the N439K viral genotype was associated with similar clinical outcomes compared to the N439 genotype (posterior mean: 0.06, 95% CI: -1.21, 1.33). **(D)** Growth curves for GLA1 (N439/D614G) or GLA2 (N439K/D614G) virus isolates in VeroE6-ACE2 cells either with or without TMPRSS2 expression. **(E)** Competition of GLA1 and GLA2 virus isolates for growth in VeroE6-ACE2 cells either with or without TMPRSS2 expression, after inoculation at a matched MOI. Quantification was by tracking the frequency of N439K within the spike gene using metagenomic NGS.

Clinical outcomes were also obtained for a subset of these patients (n=1,591), who were scored for severity of disease based on oxygen requirement: 1. No respiratory support, 2: Supplemental oxygen, 3: Invasive or non-invasive ventilation or high flow nasal cannulae, 4: Death (**Figures 4C and S2B**). Genotype counts for this analysis were N439K/D614G (n=399), D614G (with N439) (n=735) or ancestral (N439/D614) (n=457). Analysis based on our ordinal scale indicated that the N439K/D614G viral genotype was associated with similar clinical outcomes compared to D614G or ancestral genotypes (posterior mean: 0.06, 95% CI: -1.21, 1.33) (**Table S3**). All other results from the severity analysis were qualitatively similar to a previous analysis of the D614G mutation (Volz et al., 2020). These clinical data indicate that the N439K virus is not attenuated.

We next tested growth of two representative SARS-CoV-2 isolates, GLA1 (WT N439) and GLA2 (N439K), both with the D614G background (**Table S4**). Culture was carried out for 72 hours in Vero E6-ACE2 cells either with or without TMPRSS2 expression. There was no significant difference between the growth of these strains after inoculation at multiplicities of infection (MOIs) of 0.005 and 0.01. The N439K strain replicated slightly faster early after inoculation (**Figure 4D**). These data indicate that the N439K mutation does not exhibit dominant negative effects on viral growth, and most likely supports normal replication. To further assess fitness for replication in cultured cells, we carried out a cross-competition assay using inoculation of cells at a matched MOI followed by quantitation of N439 and N439K by metagenomic NGS over time (**Figure 4E**). The N439K strain demonstrated similar fitness as the WT N439 strain, with a possible fitness advantage for N439K in cells expressing TMPRSS2. Taken together with the clinical outcomes, these results indicate that the N439K mutation results in viral fitness that is similar or possibly slightly improved compared to the wild-type N439.

### The N439K variant evades antibody-mediated immunity

Having established that virus carrying the N439K mutation is fit, we sought to understand whether this mutation evades antibody-mediated immunity by evaluating recognition of the N439K variant by monoclonal antibodies and by polyclonal immune serum from 445 recovered individuals, including 6 donors who were infected by the SARS-CoV-2 N439K variant. 7.4% of the tested sera showed a greater than 2-fold reduction in binding to N439K RBD as compared to WT RBD (**Figures 5A-B and S3**). In some individuals the RBD response was diminished to low titers of <1:30 by the N439K mutation. Thus, the response to the RBD is significantly influenced by the N439K mutant within the immunodominant RBM domain (Piccoli et al., 2020) in a significant portion of persons potentially immune to WT SARS-CoV-2. The majority of sera demonstrating loss of binding were those that had overall lower responses to WT RBD, indicating lower Ab titers. The sera from the six individuals known to have recovered from infection with SARS-CoV-2 N439K virus showed no change in binding levels to WT RBD as compared to N439K RBD (**Figures 5A-B and S3**). This may reflect a true variant-specific response or that differential binding could not be measured due to the limited number of samples analyzed.

**Figure 5.**
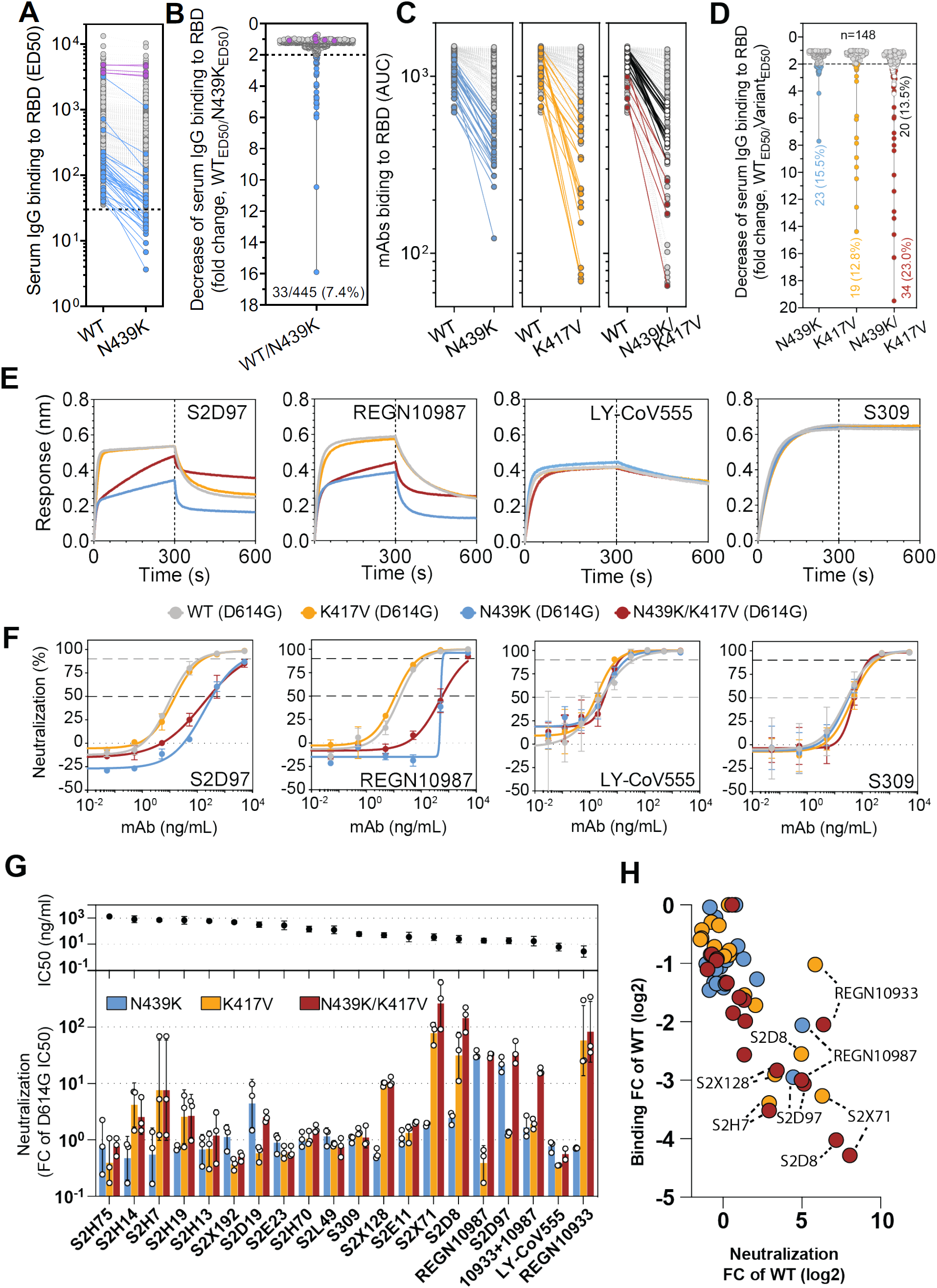
RBM variants, including N439K, exhibit immune escape from monoclonal antibodies and sera. **(A-B)** Binding of serum and plasma samples from 445 SARS-CoV-2 infected individuals against WT and N439K RBD, plotted as (A) ELISA AUC for each RBD and (B) fold change relative to WT. Blue dots indicate sera with at least 2-fold loss of binding to the N439K RBD variant as compared to WT. Purple dots indicate sera from individuals infected with SARS-CoV-2 N439K variant. **(C-D)** Binding of 144 mAbs from SARS-CoV-2 infected individuals plus four clinical-stage mAbs against WT, N439K, K417V, and N439K/K417V RBD, plotted as (C) ELISA AUC for each RBD and (D) fold change relative to WT. For all, the non-gray dots/lines indicate mAbs demonstrating at least 2-fold loss of binding to the variant RBD as compared to WT; for N439K/K417V, the mAbs corresponding to red-colored dots/lines also show a loss of binding to one of the single-site variants, whereas the mAbs corresponding to black lines/white-colored dots only lose binding to the double mutant. **(E)** Binding to RBD variants by Octet for four of the mAbs tested. **(F)** Neutralization of four VSV-pseudovirus strains by four of the mAbs tested. **(G)** Bar graph: Neutralization results for all mAbs tested, expressed as a fold-change relative to D614G (all strains are in the background of D614G). Black dots: IC50 of neutralization of D614G. **(H)** Correlation of ELISA-binding fold change and Neutralization fold change for each variant relative to WT (where a smaller ELISA AUC and therefore a smaller ratio represents loss of binding, and a larger IC50 and therefore a larger ratio represents loss of neutralization)

To understand our results at the level of individual antibodies, we evaluated a panel of 144 mAbs isolated from individuals recovered from SARS-CoV-2 infection early in the pandemic (likely with N439 WT virus) (Piccoli et al., 2020; Tortorici et al., 2020), as well as clinical-stage mAbs REGN10933, REGN10987, LY-CoV555, and S309 (the parent of VIR- 7831) (Baum et al., 2020; Hansen et al., 2020; Chen et al., 2020; Pinto et al., 2020). 15.5% of these mAbs demonstrated a >2-fold reduction of RBD binding in response to the N439K mutation (**Figures 5C-D and S4**). For comparison, we also evaluated the K417V mutation which eliminates one salt bridge at the RBM:hACE2 interface and the N439K/K417V double mutation. A similar percentage (12.8% for K417V vs 15.5% for N439K) of mAbs lost >2-fold binding to these variants, including several (13.5%) which were not sensitive to either single mutant but were sensitive to the double mutant (**Figures 5C-D**). The reduced binding of mAbs to these RBD mutants were also confirmed by bio-layer interferometry analysis (BLI) (**Figures 5E and S5A**).

To define the potential biological importance of these mutations for evasion of antibody-mediated neutralization, we tested mAbs against pseudoviruses expressing S variants N439K, K417V or N439K/K417V (**Figures 5F-H and S5B**). Neutralization of pseudoviruses containing these mutations was significantly diminished for certain mAbs, including some that are in clinical development. As predicted by its non-RBM epitope (Pinto et al., 2020), S309 was capable of neutralizing each of these variants. Sensitivity of some neutralizing mAbs to mutations at these positions have also been reported in other studies (Baum et al., 2020; Greaney et al., 2020; Li et al., 2020a; Weisblum et al., 2020) but combinations of mutations have not typically been evaluated. Overall, our results demonstrate that mutations compatible with viral fitness can result in immune evasion from both monoclonal and polyclonal antibody responses.

## DISCUSSION

The evolution of the SARS-CoV-2 RBM, a critical epitope for vaccine response and therapeutic mAbs, will depend on the fitness of RBM variants. The findings herein describe an example of a naturally-occurring RBM variant which can evade antibody-mediated immunity while maintaining fitness. Fitness of this variant, N439K, was demonstrated by repeated emergence by convergent evolution, spread to multiple countries and significant representation in the SARS-CoV-2 sequence databases, the fact that the N439K RBD retains a high affinity interaction with the hACE2 receptor, efficient viral replication in cultured cells, and no disease attenuation in a large cohort of infected individuals.

The fitness of N439K is consistent with our findings that the RBM is the most divergent region of S. This divergence indicates an ability of SARS-CoV-2 to accommodate mutations at the RBM while retaining the functional requirement of hACE2 binding, and is likely to be linked to immune pressure from neutralizing Ab responses. There is precedent for the most immunogenic region of a viral surface protein to be the fastest mutating despite harboring the receptor binding site; for example, the immunogenic globular head domain of the influenza virus hemagglutinin surface protein, which contains the sialic acid receptor binding site, evolves faster than the stalk region (Doud et al., 2018; Kirkpatrick et al., 2018). The ability to accommodate mutations in the RBM indicates a high likelihood that immune-evading SARS-CoV-2 variants compatible with fitness will continue to emerge, with implications for reinfection, vaccines, and both monoclonal and polyclonal antibody therapeutics.

In our profile of immune escape from the N439K variant, we observed resistance to a mAb currently being evaluated in clinical trials as part of a two-mAb cocktail. The promise of using cocktails of mAbs is that they should significantly lower the likelihood of drug-induced selection of resistant viruses (Baum et al., 2020). However, if circulating viral strains already carry resistant mutations to one antibody in the cocktail, this could reduce the cocktail to a monotherapy. Additionally, considering the high level of plasticity of the RBM demonstrated in the present study, there could be many combinations of RBM mutations compatible with viral fitness while leading to immune escape. This is supported by our result that N439K can compensate for a mutation (K417V) that otherwise decreases receptor binding affinity (**Figure 3D**). This particular combination of mutations is plausibly compatible with fitness as it parallels SARS-CoV RBM:hACE2 interactions (salt bridge at SARS-CoV RBD position R426 and no salt bridge at V404, **Figure 3A**). Notably, several mAbs which were not sensitive to these mutations individually were sensitive to them in combination, including the two-mAb cocktail (**Figure 5C-H**).

We propose two approaches that will be critical for minimizing the impact of mAb escape mutations. One is to develop mAbs with epitopes that are highly resistant to viral escape. This may include epitopes outside of the RBM and/or epitopes that are cross-reactive across SARS-CoV and SARS-CoV-2, indicating conserved epitopes with a low tolerance for mutation (Pinto et al., 2020; Wec et al., 2020; Wrapp et al., 2020a). A comparison of epitopes of RBM-targeting mAbs with the most conserved regions of the RBM (**Figure 1C**) may also identify RBM mAbs with a higher barrier to escape. The second approach is to screen patients, likely at the population level, for the presence of potential resistance variants prior to drug administration. The availability of multiple different mAb therapeutics in the clinic could provide the opportunity to tailor the choice of therapeutic to local circulating variants.

In general, given that access to therapeutic monoclonal antibodies via clinical trials and emergency use authorization is expanding, and as more people develop immune responses to the wildtype virus, monitoring the evolution of SARS-CoV-2 will be increasingly critical. Although SARS-CoV-2 is evolving slowly and at present should be controllable by a single vaccine (Dearlove et al., 2020), variation accumulating in the RBM could put this at risk, especially for individuals with a moderate Ab response to vaccination or infection. While we only report on evasion of antibody-mediated immunity here, it would be surprising to us if similar changes are not observed to evade T cell immunity and innate immunity.

## Acknowledgments

We thank all Scottish NHS virology laboratories who provided samples for sequencing and Scott Arkison for HPC maintenance. We thank Chiara Silacci-Fregni from Humabs BioMed, Sandra Jovic, Blanca Fernandez Rodriguez, Federico Mele, from the Institute for Research in Biomedicine in Bellinzona and Tatiana Terrot from Ente Ospedaliero Cantonale in Lugano for the help in collecting sera samples. We thank Cindy Ng for help with protein production. We thank Julia Di Iulio for help with analyzing GISAID sequences. We gratefully acknowledge the authors, originating and submitting laboratories of the sequences from GISAID, https://www.gisaid.org, on which much of this research is based.

The ISARIC WHO CCP-UK study protocol is available at https://isaric4c.net/protocols; study registry https://www.isrctn.com/ISRCTN66726260. This work uses data provided by patients and collected by the NHS as part of their care and support #DataSavesLives. We are grateful to the 2648 frontline NHS clinical and research staff and volunteer medical students who collected the data in challenging circumstances; and the generosity of the participants and their families for their individual contributions in these difficult times. We also acknowledge the support of Jeremy J Farrar, Nahoko Shindo, Devika Dixit, Nipunie Rajapakse, Lyndsey Castle, Martha Buckley, Debbie Malden, Katherine Newell, Kwame O’Neill, Emmanuelle Denis, Claire Petersen, Scott Mullaney, Sue MacFarlane, Nicole Maziere, Julien Martinez, Oslem Dincarslan, and Annette Lake.

For funding, we thank: MRC (MC UU 1201412), Wellcome Trust Collaborator Award (206298/Z/17/Z – ARTIC Network; TCW Wellcome Trust Award 204802/Z/16/Z and Chief Scientist Office Project (COV/EDI/20/11). COG-UK is supported by funding from the Medical Research Council (MRC) part of UK Research & Innovation (UKRI), the National Institute of Health Research (NIHR) and Genome Research Limited, operating as the Wellcome Sanger Institute. National Institute for Health Research (NIHR; award CO-CIN- 01), the Medical Research Council (MRC; grant MC_PC_19059), and by the NIHR Health Protection Research Unit (HPRU) in Emerging and Zoonotic Infections at University of Liverpool in partnership with Public Health England (PHE), in collaboration with Liverpool School of Tropical Medicine and the University of Oxford (award 200907), NIHR HPRU in Respiratory Infections at Imperial College London with PHE (award 200927), Wellcome Trust and Department for International Development (DID; 215091/Z/18/Z), the Bill and Melinda Gates Foundation (OPP1209135), Liverpool Experimental Cancer Medicine Centre (grant reference C18616/A25153), NIHR Biomedical Research Centre at Imperial College London (IS-BRC-1215-20013), EU Platform for European Preparedness Against (Re-)emerging Epidemics (PREPARE; FP7 project 602525), and NIHR Clinical Research Network for providing infrastructure support for this research. PJMO is supported by a NIHR senior investigator award (201385). The views expressed are those of the authors and not necessarily those of the Department of Health and Social Care, DID, NIHR, MRC, Wellcome Trust, or PHE.

## Author Contributions

Conceived research: E.C.T., R. Sp., H.W.V., A.T., D.C., D.L.R., G.S.

Designed experiments: E.C.T., A.T., D.C., G.S.

Donors’ Recruitment and Sample Collection for serological analysis and mAbs isolation: A.C., P.F., F.S., C.G., M.G., A.Ri., A.H., M.G.S., P.J.M.O., J.K.B.

Isolation of mAbs: D.P., K.C., F.Z., M.B., M.P., E.C.

Expressed and purified proteins: J.D., N.C., M.M., S.J.

Performed binding and neutralization assays: L.E.R., J.A.W., L.P., A.D.M., J.B., S.J. Performed NGS sequencing and analysis: E.C.T., A.S.F., J.H., V.B.S., K.N., L.T., N.J., D.M., K.S., S.C., J.Nic.

Performed phylogenetic and epidemiological analysis: S.L., A.O., R.M.C., B.J., A.Ra., J.H., S.J.L., D.L.R.

Performed cross-competition and growth assays of primary isolates: A.S.F., C.D., A.W., S.J.R.

Collected and analyzed clinical data: E.C.T., J.G.S., D.J.P., R. Sh., N.J., K.L. Performed and analyzed real-time PCR assays: R. Sh., N.J., R.F.J.

Analyzed data: E.C.T., L.E.R., J.G.S., R. Sp., J.A.W., L.P., D.J.P., A.Ra., J.Nix, S.J.L., D.L.R., G.S.

Wrote the manuscript: E.C.T., L.E.R., H.W.V., A.T., D.C., D.L.R., G.S.

Supervised the project: E.C.T., D.L.R., G.S.

## DECLARATION OF INTERESTS

L.E.R., R. Sp., J.A.W., L.P., J.D., N.C., M.M., A.D.M., J.B., D.P., K.C., F.Z., S.J., M.B., M.P., E.C., H.W.V., A.T., D.C., and G.S. are or were employees of Vir Biotechnology Inc. and may hold shares in Vir Biotechnology Inc. C.G. is a consultant to Humabs BioMed SA. J.Nix is a consultant with Vir Biotechnology Inc. The other authors declare no competing interests.

## MATERIALS AND METHODS

### Sample donors

Samples from 439 SARS-CoV-2 infected individuals were obtained from the Ticino healthcare workers cohort (Switzerland), described previously (Piccoli et al., 2020), and under study protocols approved by the local Institutional Review Board (Canton Ticino Ethics Committee, Switzerland). All donors provided written informed consent for the use of blood and blood components (such as PBMCs, sera or plasma). In the Ticino region of Switzerland and during the time period of collection (February-March 2020) no N439K SARS-CoV-2 isolates were reported.

Samples from six N439K variant infected individuals were obtained from the ISARIC4C consortium (https://isaric4c.net/). Ethical approval was given by the South Central-Oxford C Research Ethics Committee in England (reference 13/SC/0149), and by the Scotland A Research Ethics Committee (reference 20/SS/0028). The study was registered at https://www.isrctn.com/ISRCTN66726260.

Residual nucleic acid extracts derived from the nose-throat swabs of 1918 SARS-CoV-2 positive individuals whose diagnostic samples were submitted to the West of Scotland specialist virology centre between 3^rd^ March and 30^th^ June 2020 were sequenced as part of the COG-UK consortium under study protocols approved by the relevant national biorepositories (16/WS/0207NHS and 10/S1402/33) (, 2020).

### Structural analysis

RBM residues were determined based on the RBD:ACE2 complex crystal structures 2AJF for SARS-CoV (Li et al., 2005) and 6M0J for SARS-CoV-2 (Lan et al., 2020). The 2AJF structure was obtained from the PDB-REDO server (pdb-redo.eu) and was subsequently prepared in the molecular modeling software MOE (v2019.0102, https://www.chemcomp.com) using the structure preparation, protonation and energy minimization steps with default settings. RBD residues within 6.0A distance of any ACE2 atoms (determined using MOE) were determined for each of the two copies of the complex in the asymmetric unit, and then were combined to obtain the RBM. 6M0J was obtained from the Coronavirus Structural Task Force server (https://github.com/thorn-lab/coronavirus_structural_task_force) and was further refined (using Refmac5 v5.8.0258), manually fitted (using Coot v0.9) and prepared (using MOE, as described above) in multiple iterative cycles. The final structure was analyzed for RBD-ACE2 contact residues with a 6.0A cutoff to obtain the RBM (using MOE). The final list of RBM residues (**Figure 1C**) was arrived at by combining the SARS-CoV and SARS-CoV-2 results.

Using MOE, the pairwise binding energy between each residue in SARS-CoV-2 RBD and each residue in ACE2, and the total binding energy for all interactions, was determined at cutoff distances 3.0A, 3.5A, 4.0A, 4.5A, 5.0A, 5.5A, 6.0A, 6.5A and 7.0A. The percentage of the total binding energy for each interacting RBD residue was calculated for each distance cutoff and was then averaged over all cutoffs. The resulting values are shown in green in **Figure 1C**.

### Determining conservation from deposited GISAID sequences

Differential accumulation of amino acid variants in the RBM, RBD or Spike protein was computed taking into account only the presence or absence of a variant at any residue. Each variant called present counts one. A variant is called present if there are at least X number of supporting sequences deposited in GISAID, where X varies from 2 to 20. The number of variants is then normalized to the size of the domain (number of residues).

### Evaluation of deep mutational scanning (DMS) data

DMS data was retrieved from (Starr et al., 2020). Variant-level DMS scores were aggregated to residue-level by taking the minimum (most disruptive variant) or the average score across all variants of a residue, except for the reference residue and the stop codon. Alternatively, minimum and average scores are computed only across variants that have been observed as naturally occurring. Data were represented as a heatmap annotated with: frequency of non-reference amino acids in deposited GISAID sequences (n ≈ 130,000, at least 4 sequences were required to call a variant as present), in Log10 scale; number of countries in which a variant was observed; and percentage of total binding energy computed from an X-ray crystal structure (cf. structural analysis methods section).

### Recombinant glycoprotein production

Prefusion-stabilized SARS-CoV-2 spike protein variants (residues 14-1211, containing the 2P and Furin cleavage site mutations (Walls et al., 2020) with a mu-phosphatase signal sequence and a C-terminal Avi-8xHis-EPEA-tag in a pD2610-V5 vector (ATUM Bio) were expressed in Expi293F cells at 37°C and 8% CO_2_ according to manufacturer’s instructions (Thermo Fisher Scientific). Cell culture supernatant was collected after four days and purified over a 5 mL C-tag affinity matrix (Thermo Fisher Scientific). Elution fractions were concentrated and injected on a Superose 6 Increase 10/300 GL column with 1x PBS pH 7.4 as running buffer.

SARS-CoV-2 RBD variants (residues 328-531 with a C-terminal thrombin-cleavage site-TwinStrep-8xHis-tag, and N-terminal signal sequence) were expressed in Expi293F cells at 37°C and 8% CO_2_ in a humidified incubator. Transfection was performed using ExpiFectamine 293 reagent (Thermo Fisher Scientific). Cell culture supernatant was collected three days after transfection and supplemented with 10x PBS to a final concentration of 2.5x PBS (342.5 mM NaCl, 6.75 mM KCl and 29.75 mM phosphates), or 3.2x for RBD N439R. SARS-CoV-2 RBDs were purified using 1 or 5 ml HisTALON superflow cartridges (Takara Bio) and subsequently buffer exchanged into Cytiva 1x HBS-N buffer or PBS.

RBDs from other sarbecoviruses were expressed in Expi293F cells at 37°C and 8% CO_2_. Cells were transfected using PEI MAX. Cell culture supernatant was collected seven days after transfection. Proteins were purified using a 5 ml Strep-Tactin XT Superflow high capacity cartridge followed by buffer exchange to PBS using HiPrep 26/10 desalting columns.

For S binding measurements, recombinant ACE2 (residues 19-615 from Uniprot Q9BYF1 with a C-terminal thrombin cleavage site-TwinStrep-10xHis-GGG-tag, and N-terminal signal sequence) was expressed in Expi293 cells at 37°C and 8% CO_2_ in a humified incubator. Transfection was performed using ExpiFectamine 293 reagent (Thermo Fisher Scientific). Cell culture supernatant was collected seven days after transfection, supplemented with buffer to a final concentration of 80 mM Tris-HCl pH 8.0, 100 mM NaCl, and then incubated with BioLock solution for one hour. After filtration through a 0.22 µm filter, ACE2 was purified using a 1 ml StrepTrap HP column (Cytiva) followed by isolation of the monomeric ACE2 by size exclusion chromatography using a Superdex 200 Increase 10/300 GL column pre-equilibrated in PBS (Gibco 10010-023).

For binding measurements with surface-captured RBD, recombinant ACE2 (residues 19-615 from Uniprot Q9BYF1 with a C-terminal AviTag-10xHis-GGG-tag, and N-terminal signal sequence) was expressed in HEK293.sus using standard methods (ATUM Bio). Protein was purified via Ni Sepharose resin followed by isolation of the monomeric ACE2 by size exclusion chromatography using a Superdex 200 Increase 10/300 GL column pre-equilibrated with PBS.

For binding measurements with surface-captured ACE2, recombinant ACE2 (residues 18-615 with a C-terminal GS-IgG2a-Mm-Fc tag, and N-terminal signal sequence) was stably transfected in CHO-K1 GS knock-down cell line (ATUM Bio). Protein was purified via protein A and buffer exchanged into PBS.

### Binding measurements using surface plasmon resonance (SPR)

SPR binding measurements were performed using a Biacore T200 instrument. S protein was surface captured via anti-AviTag pAb covalently immobilized on a CM5 chip, RBD protein was surface captured via StrepTactin XT covalently immobilized on a CM5 chip, and ACE2-mFc was surface captured via covalent immobilization of the Cytiva Mouse antibody capture kit on a C1 chip. Running buffer was Cytiva HBS-EP+ (pH 7.4) and all measurements were performed at 25 °C. All experiments were performed as single-cycle kinetics, with a 3-fold dilution series of monomeric ACE2 starting from 300 nM, each concentration injected for 180 sec, or a 3-fold dilution series of RBD starting from 50 nM, each concentration injected for 240 sec. All data were double reference-subtracted and fit to a binding model using Biacore Evaluation software. For one representative replicate, capture levels were normalized to WT for visualization. Binding data with ACE2 as analyte were fit to a 1:1 binding model. Binding data with RBD as analyte were fit to a Heterogeneous Ligand binding model, due to an artifactual kinetic phase with very slow dissociation that arises when RBD is an analyte; the lower affinity of the two K_D_s reported by the fit is reported as the K_D_ of the RBD-ACE2 interaction (the two reported K_D_s are separated by at least two orders of magnitude for all fits). The measured K_D_ for ACE2 binding to S is likely influenced by conformational dynamics of the RBDs in the context of the prefusion S trimer. Reported K_D_s are an average of 3-4 replicates measured on at least two separate days, with error given as SEM.

### Epidemiological and genome surveillance

A national sequencing collaboration formed at the start of the epidemic in the UK, CoG-UK consortium (, 2020), has facilitated the tracking of SARS-CoV-2 sequences across Scotland since the start of the outbreak in February 2020 (6,825 sequences by Oct 6, 2020) and real-time monitoring of genetic changes in the Spike gene that might be associated with changes in virulence or transmissibility. Sequencing was carried out using an amplicon-based protocol in real-time at a rate of up to 300 genomes per week. 50% of samples were selected as surveillance samples, representing Scottish health boards proportionately based on population size, while 50% were selected to allow intervention with local issues such as nosocomial infection in hospitals and nursing homes. A gradual increase in the prevalence of the N439K polymorphism was noted to become increasingly prevalent during April 2020. This was noted to be particularly common in the Greater Glasgow & Clyde NHS health board region but spread to adjacent Scottish health boards also.

Sequencing libraries were prepared according to the ARTIC nCoV-2019 described in detail at https://artic.network/ncov-2019. Briefly, PCR amplicons were generated using the nCoV-2019 PrimalSeq sequencing primers using 25-35 cycles of amplification. Generated amplicons were used to prepare either Oxford Nanopore or Illumina sequencing libraries. Oxford Nanopore libraries were prepared as described in the link above and sequenced in a flow cell R9.4.1 (Oxford Nanopore Technologies, Part Number FLO- MIN106D), using MinKNOW version 19.12.6. Raw FAST5 files were basecalled using Guppy version 3.2.10 in high accuracy mode with a minimum quality score of 7. Reads were size filtered, demultiplexed and trimmed with Porechop (https://github.com/rrwick/Porechop), and mapped against reference strain Wuhan-Hu-1 (MN908947). Variants were called using Nanopolish 0.11.3 and accepted if they had a log-likelihood score of greater than 200 and minimum read coverage of 20. For Illumina sequencing, amplicons were used to prepare libraries using the Kapa HyperPrep kit (Kapa Biosystems, Part Number KK8504) and further processed as described in the competition assay sequencing method. Sequencing was carried out on Illumina’s MiSeq system (Illumina, Part Number SY-410-1003) using a MiSeq Reagent v2 500 cycle kit (Illumina, Part Number MS-102-2003). Reads were trimmed with trim_galore (http://www.bioinformatics.babraham.ac.uk/projects/trim_galore/) and mapped with BWA (Li and Durbin, 2009)) to the Wuhan-Hu-1 (MN908947) reference sequence, followed by primer trimming and consensus calling with iVar (Grubaugh et al., 2019) and a minimum read coverage of 10.

### Phylogenetic and phylodynamic analysis

UK sequences were obtained from the COG-UK consortium, https://www.cogconsortium.uk. Global sequences were obtained from the GISAID Initiative, https://www.gisaid.org on Oct 19 2020. The sequences were mapped using minimap2 and padded against the Wuhan/WH04/2020 reference. The sequences were downsampled with weights that normalise sequence count per epiweek, maximise the number of countries and lineages represented, and enriching for sequences with the N439K mutation. A maximum-likelihood phylogenetic tree was constructed using IQ-TREE with the the following parameters: -czb -blmin 0.0000000001 -m HKY --runs 5 and all other parameters set to default. The tree was visualised with custom python code using the baltic library, https://github.com/evogytis/baltic.

For the phylodynamic analysis, Scottish “introduction” lineages were identified (Lycett et al., 2020, in prep and see http://sars2.cvr.gla.ac.uk/RiseFallScotCOVID), and the skygrowth package in R was used to estimate the effective population size over time, and the growth rate of the lineage within Scotland (Volz and Frost, 2017).

### Evaluation of clinical samples

Clinical samples submitted to the West of Scotland Specialist Virology Centre for SARS-CoV-2 diagnostic rt-PCR testing were selected for sequencing as part of the COVID-19 UK Genomics UK Consortium (COG-UK) project, resulting in 1918 whole genome sequences originating from the NHS Greater Glasgow and Clyde Health Board region. Sequences were linked to electronic patient records and basic metadata including sample date, age, sex, admission to hospital and mortality at 28 days post diagnosis extracted. The electronic patient records of a subset of 1591 patients underwent full case-note review and clinical severity was recorded based on a 4-level ordinal scale: 1. no requirement for respiratory support, 2. treatment with supplemental oxygen via facemask or low-flow nasal cannulae, 3. intubation and ventilation, non-invasive ventilation or oxygen delivery by high flow nasal cannulae devices, 4. death within the 28 days following diagnosis. We modified the WHO ordinal scale to these 4 points as described previously (Volz et al., 2020) to avoid using hospitalisation as a criterion of severity because 1) many patients in nursing homes had severe infection but were not admitted to hospital, and 2) early in the outbreak, all cases were hospitalised irrespective of the severity of their infection.

These data had previously been analysed to test for an effect of the D614G mutation on the severity of disease (Volz et al., 2020); we extend that analysis here using the same methodology to test for an effect of the N439K mutation. Additionally, we perform a new analysis using a model with the same structure to test for an effect of both the D614G mutation and the D614G/N439K mutation combination on the viral load of infected patients, as measured by cycle threshold value. In both cases we cannot estimate the marginal effect of the N439K mutation, as we only have the mutation on the 614G genetic background, so the individual effect of N439K cannot be separated from any potential epistatic interactions between the mutations.

Briefly, the structure of the model used previously (Volz et al., 2020) and in the present study is a phylogenetic generalised additive model with mutation being the primary predictor of interest. The model controls for biological sex, age and the number of days since the first reported case in the dataset, with the latter two being included as penalised splines with a maximum of 30 knots. If the patient was part of a cluster of cases, this was included as a random effect, with individuals not part of clusters being assigned their own levels. Correlations driven by the rest of the genome are controlled for by a phylogenetic random effect using a correlation matrix generated under a Brownian motion assumption from a phylogeny estimated in IQ-TREE 2 v. 2.0.6 (Minh et al., 2020) using a HKY + Γ model, masking the positions recommended by De Maio et al. as of 22/7/2020 (https://virological.org/t/issues-with-sars-cov-2-sequencing-data/473/13), rooted on the first sequenced SARS-CoV-2 genome (Wu et al., 2020). The priors for the severity model were those used in the previous analysis of this data. The priors for the model of the viral load were a student-t (mean = 20, scale = 10, degrees of freedom = 3) prior on the model intercept, a Gaussian (mean = 0, standard deviation = 10) prior over the fixed effects, and an exponential (lambda = 0.1) prior over the random effect, penalised spline and residual standard deviations.

There are two key structural differences between the model used previously (Volz et al., 2020) and the model used here. Firstly, mutation is a three level rather than two level factor (D614/N439, D614G/N439 and D614G/N439K) with the ancestral D614/N439 being the reference level. Secondly, as we are now interested in two mutations, we estimated the phylogeny used to control for the effect of the rest of the genome excluding both the nucleotide position underlying the D614G mutation and the nucleotide position underlying the N439K mutation (in addition to the sites from De Maio et al mentioned above).

The severity model used a cumulative error structure while the model on the CT values used a Gaussian error structure. In both cases, the models were estimated in brms v. 2.13.5 (Bürkner, 2018). The presented models had no divergent transitions, Rhat values less than 1.01, and appropriate bulk and tail effective sample sizes for all parameters. Shortest probability intervals were calculated using the R package SPIn v. 1.1 (Liu et al., 2015). Analysis code is available at https://github.com/dpascall/SARS-CoV-2-mutation-analysis.

### qPCR of clinical samples

All samples were tested in duplicate using the 2019-nCoV_N1 assay RT-qPCR assay (https://www.fda.gov/media/134922/download). Ready-mixed primers and probe were obtained from IDT (Leuven, Belgium). PCR was carried out using NEB Luna Universal Probe One-Step Reaction Mix and Enzyme Mix (New England Biolabs, Herts, UK), primers and probe at 500 nM and 127.5 nM, respectively, and 5 µl of RNA sample in a final volume of 20 µl. No template negative controls were included after every seventh sample. Six ten-fold dilutions of SARS-CoV-2 RNA standards were tested in duplicate in each assay; standards were calibrated using a plasmid containing the N sequence that had been quantified using droplet digital PCR. Thermal cycling was performed on an Applied Biosystems™ 7500 Fast PCR instrument running SDS software v2.3 (ThermoFisher Scientific) under the following conditions: 55 °C for 10 minutes and 95 °C for 1 minute followed by 45 cycles of 95 °C for 10 s and 58 °C for 1 minute. Assays were repeated if the reaction efficiency was <90% or the R2 value of the standard curve was ≤0.998. Where possible, testing of samples was repeated if the %CV of the duplicates was <10%.

### Viral growth curve

VeroE6-ACE2 cells (VeroE6 cells induced to overexpress Ace2) either with or without TMPRSS2 overexpression (Rhin et al., 2020 under review) were seeded in a 12- well plate and inoculated with an MOI of 0.01 with either the GLA1 (N439/D614G) or GLA2 (N439K/D614G) virus isolates for 1hr before washing the cells three times in PBS and replacing with 2% DMEM. 100ul of media was removed at each timepoint, RNA was extracted, and the presence of SARS-CoV-2 determined using 2019-nCOV-N1 assays (IDT) with an NEB Luna Universal Probe One-Step RT-qPCR Kit. A standard curve was used to determine the copy number present per ml of cell culture media. 100ul of the fresh media was also tested for the presence of virus, which was undetectable in all wells.

### Competition assay

Three T25 Flasks were seeded with VeroE6-Ace2 or VeroE6-Ace2-TMPRSS2 and inoculated with either single viruses or both GLA1 and GLA2 virus strains at an MOI of 0.01 for 1 hr. The flasks were washed three times with PBS, with 100 ul of the final wash being retained to determine the presence of free virus, before adding 5 ml of fresh 2% DMEM. At 24, 48, and 72 hrs, 500 ul of media was removed, which was replaced with 500 ul fresh media. 300 ul was used for RNA extraction and NGS analysis of the frequencies of the specific positions within the spike protein. The single virus inoculations showed no alternations in the frequency of the amino acid positions and the final wash showing no free virus in the supernatant. We used an unbiased metagenomic NGS sequencing pipeline to quantify variation across the whole viral genome on the Illumina NGS Next Seq platform. Briefly, extracted nucleic acid was incubated with DNaseI (Thermo Fisher, Part Number AM2222) followed by cDNA synthesis using SuperScript III (Thermo Scientific, Part Number 18080044) and NEBNext Ultra II Non-Directional RNA Second Strand Synthesis Module (New England Biolabs, Part Number E6111L). Samples were further processed using the Kapa LTP Library Preparation Kit for Illumina Platforms (Kapa Biosystems, Part Number KK8232) and indexed with the NEBNext Multiplex Oligos for Illumina 96 Unique Dual Index Primer Pairs (New England Biolabs, Part Number E6442S). Libraries were sequenced on Illumina’s NextSeq 550 System (Illumina, Part Number SY- 415-1002), generating 10 million pairs of reads per sample.

### Ab discovery and recombinant expression

Human mAbs were isolated from plasma cells or memory B cells of SARS-CoV or SARS-CoV-2 immune donors, as previously described (Corti et al., 2011; Pinto et al., 2020; Tortorici et al., 2020). LY-CoV555 mAb was obtained from Eli Lilly and Company. REGN10933 and REGN10987 mAbs were produced recombinantly based on published sequences (Hansen et al., 2020).

### Enzyme-linked immunosorbent assay (ELISA)

A total of 148 human monoclonal antibodies or 445 human sera were tested for binding to RBD WT and mutants. Spectraplate-384 plates with high protein binding treatment (custom made from Perkin Elmer) were coated overnight at 4 °C with 0.5 µg/ml (for mAbs) or 5 ug/ml (for sera) SARS-CoV-2 RBD WT, N439K, K417V or N439K/K417V in phosphate-buffered saline (PBS), pH 7.2. Plates were subsequently blocked with Blocker Casein 1% supplemented with 0.05% Tween 20 (Sigma-Aldrich) for 1 h at room temperature. The coated plates were incubated with serial dilutions of the monoclonal antibodies or of the sera for 1 h at room temperature. The plates were then washed with PBS containing 0.1% Tween-20 (PBS-T), and alkaline phosphatase-goat anti-human IgG (Southern Biotech) was added and incubated for 1 h at room temperature. After 3 washing steps with PBS-T, P-NitroPhenyl Phosphate (pNPP, Sigma-Aldrich) substrate was added and incubated for 30 min at room temperature. The absorbance of 405 nm was measured by a microplate reader (Biotek). Fitting was performed using a 4-parameter logistic (4PL) model, yielding dose-response curves from which the area under the curve (AUC) between 5 and 500 ng/ml was computed. The AUC allows to capture, in a single metric, shifts of interest in two parameters of the 4PL model: EC50 and upper asymptote.

### Binding measurements using BLI

BLI binding measurement was performed on a selection of human monoclonal antibodies tested by ELISA. Antibodies were diluted to 2.7 µg/ml in kinetic buffer (PBS supplemented with 0.05% BSA) and immobilized on Protein A Biosensors of an Octet RED96 system (FortéBio). Antibody-coated biosensors were incubated for 5 min with a solution containing 5 µg /ml of SARS-CoV2 RBD WT, N439K, K417V or N439/K417V in kinetic buffer. A dissociation step was then performed by incubating the biosensors for 5 min in kinetic buffer. Change in molecules bound to the biosensors caused a shift in the interference pattern that was recorded in real time and plotted using GraphPad Prism 8 software.

### VSV Pseudovirus generation

Replication defective VSV pseudovirus (Takada et al., 1997) expressing SARS-CoV-2 spike protein were generated as previously described (Riblett et al., 2016) with some modifications. Plasmids encoding SARS-CoV-2 spike variants were generated by site-directed mutagenesis of the wild-type plasmid, pcDNA3.1(+)-spike-D19 (Giroglou et al., 2004). Lenti-X™ 293T cells (Takara, 632180) were seeded in 10-cm dishes at a density of 1e5 cells/cm^2^ and the following day transfected with 5 μg of spike expression plasmid with TransIT-Lenti (Mirus, 6600) according to the manufacturer’s instructions. One day post-transfection, cells were infected with VSV-luc (VSV-G) (Kerafast, EH1020-PM) for 1 h, rinsed three times with PBS, then incubated for an additional 24 h in complete media at 37°C. The cell supernatant was clarified by centrifugation, filtered (0.45 μm), aliquoted, and frozen at -80°C.

### Pseudovirus neutralizations

Vero E6 cells (ATCC CRL-1586) were seeded into clear bottom white 96 well plates (Costar, 3903) at a density of 2e4 cells per well. The next day, mAbs were serially diluted in pre-warmed complete media, mixed at a 1:1 ratio with pseudovirus and incubated for 1 h at 37°C in round bottom polypropylene plates. Media from cells was aspirated and 50 μL of virus-mAb complexes were added to cells and then incubated for 1 h at 37°C. An additional 100 μL of prewarmed complete media was then added on top of complexes and cells incubated for an additional 16-24 h. Conditions were tested in duplicate wells on each plate and at least six wells per plate contained uninfected, untreated cells (mock) and infected, untreated cells (‘no mAb control’). Virus-mAb-containing media was then aspirated from cells and 100 mL of a 1:4 dilution of Bio-glo (Promega, G7940) in PBS was added to cells. Plates were incubated for 10 mins at room temperature and then were analyzed on the Envision plate reader (PerkinElmer). Relative light units (RLUs) for infected wells were subtracted by the average of RLU values for the mock wells (background subtraction) and then normalized to the average of background subtracted “no mAb control” RLU values within each plate. Percent neutralization was calculated by subtracting from 1 the normalized mAb infection condition. Data were analyzed and visualized with Prism (Version 8.4.3). IC50 curves were calculated from the interpolated value from the log(inhibitor) vs. response – variable slope (four parameters) nonlinear regression with an upper constraint of <100. Each neutralization infection was conducted on three independent days.

## APPENDICES

### Appendix 1: ISARIC4C Investigators

Consortium Lead Investigator: J Kenneth Baillie, Chief Investigator: Malcolm G Semple, Co-Lead Investigator: Peter JM Openshaw. ISARIC Clinical Coordinator: Gail Carson. Co-Investigators: Beatrice Alex, Benjamin Bach, Wendy S Barclay, Debby Bogaert, Meera Chand, Graham S Cooke, Annemarie B Docherty, Jake Dunning, Ana da Silva Filipe, Tom Fletcher, Christopher A Green, Ewen M Harrison, Julian A Hiscox, Antonia Ying Wai Ho, Peter W Horby, Samreen Ijaz, Saye Khoo, Paul Klenerman, Andrew Law, Wei Shen Lim, Alexander J Mentzer, Laura Merson, Alison M Meynert, Mahdad Noursadeghi, Shona C Moore, Massimo Palmarini, William A Paxton, Georgios Pollakis, Nicholas Price, Andrew Rambaut, David L Robertson, Clark D Russell, Vanessa Sancho-Shimizu, Janet T Scott, Thushan de Silva, Louise Sigfrid, Tom Solomon, Shiranee Sriskandan, David Stuart, Charlotte Summers, Richard S Tedder, Emma C Thomson, AA Roger Thompson, Ryan S Thwaites, Lance CW Turtle, Maria Zambon. Project Managers: Hayley Hardwick, Chloe Donohue, Ruth Lyons, Fiona Griffiths, Wilna Oosthuyzen. Data Analysts: Lisa Norman, Riinu Pius, Tom M Drake, Cameron J Fairfield, Stephen Knight, Kenneth A Mclean, Derek Murphy, Catherine A Shaw. Data and Information System Managers: Jo Dalton, James Lee, Daniel Plotkin, Michelle Girvan, Egle Saviciute, Stephanie Roberts, Janet Harrison, Laura Marsh, Marie Connor, Sophie Halpin, Clare Jackson, Carrol Gamble. Data integration and presentation: Gary Leeming, Andrew Law, Murray Wham, Sara Clohisey, Ross Hendry, James Scott-Brown. Material Management: William Greenhalf, Victoria Shaw, Sarah McDonald. Patient engagement: Seán Keating Outbreak Laboratory Staff and Volunteers: Katie A. Ahmed, Jane A Armstrong, Milton Ashworth, Innocent G Asiimwe, Siddharth Bakshi, Samantha L Barlow, Laura Booth, Benjamin Brennan, Katie Bullock, Benjamin WA Catterall, Jordan J Clark, Emily A Clarke, Sarah Cole, Louise Cooper, Helen Cox, Christopher Davis, Oslem Dincarslan, Chris Dunn, Philip Dyer, Angela Elliott, Anthony Evans, Lorna Finch, Lewis WS Fisher, Terry Foster, Isabel Garcia-Dorival, Willliam Greenhalf, Philip Gunning, Catherine Hartley, Antonia Ho, Rebecca L Jensen, Christopher B Jones, Trevor R Jones, Shadia Khandaker, Katharine King, Robyn T. Kiy, Chrysa Koukorava, Annette Lake, Suzannah Lant, Diane Latawiec, L Lavelle-Langham, Daniella Lefteri, Lauren Lett, Lucia A Livoti, Maria Mancini, Sarah McDonald, Laurence McEvoy, John McLauchlan, Soeren Metelmann, Nahida S Miah, Joanna Middleton, Joyce Mitchell, Shona C Moore, Ellen G Murphy, Rebekah Penrice-Randal, Jack Pilgrim, Tessa Prince, Will Reynolds, P. Matthew Ridley, Debby Sales, Victoria E Shaw, Rebecca K Shears, Benjamin Small, Krishanthi S Subramaniam, Agnieska Szemiel, Aislynn Taggart, Jolanta Tanianis-Hughes, Jordan Thomas, Erwan Trochu, Libby van Tonder, Eve Wilcock, J. Eunice Zhang. Local Principal Investigators: Kayode Adeniji, Daniel Agranoff, Ken Agwuh, Dhiraj Ail, Ana Alegria, Brian Angus, Abdul Ashish, Dougal Atkinson, Shahedal Bari, Gavin Barlow, Stella Barnass, Nicholas Barrett, Christopher Bassford, David Baxter, Michael Beadsworth, Jolanta Bernatoniene, John Berridge, Nicola Best, Pieter Bothma, David Brealey, Robin Brittain-Long, Naomi Bulteel, Tom Burden, Andrew Burtenshaw, Vikki Caruth, David Chadwick, Duncan Chambler, Nigel Chee, Jenny Child, Srikanth Chukkambotla, Tom Clark, Paul Collini, Catherine Cosgrove, Jason Cupitt, Maria-Teresa Cutino-Moguel, Paul Dark, Chris Dawson, Samir Dervisevic, Phil Donnison, Sam Douthwaite, Ingrid DuRand, Ahilanadan Dushianthan, Tristan Dyer, Cariad Evans, Chi Eziefula, Chrisopher Fegan, Adam Finn, Duncan Fullerton, Sanjeev Garg, Sanjeev Garg, Atul Garg, Effrossyni Gkrania-Klotsas, Jo Godden, Arthur Goldsmith, Clive Graham, Elaine Hardy, Stuart Hartshorn, Daniel Harvey, Peter Havalda, Daniel B Hawcutt, Maria Hobrok, Luke Hodgson, Anil Hormis, Michael Jacobs, Susan Jain, Paul Jennings, Agilan Kaliappan, Vidya Kasipandian, Stephen Kegg, Michael Kelsey, Jason Kendall, Caroline Kerrison, Ian Kerslake, Oliver Koch, Gouri Koduri, George Koshy, Shondipon Laha, Steven Laird, Susan Larkin, Tamas Leiner, Patrick Lillie, James Limb, Vanessa Linnett, Jeff Little, Michael MacMahon, Emily MacNaughton, Ravish Mankregod, Huw Masson, Elijah Matovu, Katherine McCullough, Ruth McEwen, Manjula Meda, Gary Mills, Jane Minton, Mariyam Mirfenderesky, Kavya Mohandas, Quen Mok, James Moon, Elinoor Moore, Patrick Morgan, Craig Morris, Katherine Mortimore, Samuel Moses, Mbiye Mpenge, Rohinton Mulla, Michael Murphy, Megan Nagel, Thapas Nagarajan, Mark Nelson, Igor Otahal, Mark Pais, Selva Panchatsharam, Hassan Paraiso, Brij Patel, Natalie Pattison, Justin Pepperell, Mark Peters, Mandeep Phull, Stefania Pintus, Jagtur Singh Pooni, Frank Post, David Price, Rachel Prout, Nikolas Rae, Henrik Reschreiter, Tim Reynolds, Neil Richardson, Mark Roberts, Devender Roberts, Alistair Rose, Guy Rousseau, Brendan Ryan, Taranprit Saluja, Aarti Shah, Prad Shanmuga, Anil Sharma, Anna Shawcross, Jeremy Sizer, Manu Shankar-Hari, Richard Smith, Catherine Snelson, Nick Spittle, Nikki Staines, Tom Stambach, Richard Stewart, Pradeep Subudhi, Tamas Szakmany, Kate Tatham, Jo Thomas, Chris Thompson, Robert Thompson, Ascanio Tridente, Darell Tupper-Carey, Mary Twagira, Andrew Ustianowski, Nick Vallotton, Lisa Vincent-Smith, Shico Visuvanathan, Alan Vuylsteke, Sam Waddy, Rachel Wake, Andrew Walden, Ingeborg Welters, Tony Whitehouse, Paul Whittaker, Ashley Whittington, Meme Wijesinghe, Martin Williams, Lawrence Wilson, Sarah Wilson, Stephen Winchester, Martin Wiselka, Adam Wolverson, Daniel G Wooton, Andrew Workman, Bryan Yates, and Peter Young.

### Appendix 2: COG-UK

**Funding acquisition, leadership, supervision, metadata curation, project administration, samples, logistics, Sequencing, analysis, and Software and analysis tools:**

Thomas R Connor ^33, 34^, and Nicholas J Loman ^15^.

**Leadership, supervision, sequencing, analysis, funding acquisition, metadata curation, project administration, samples, logistics, and visualisation:**

Samuel C Robson ^68^.

**Leadership, supervision, project administration, visualisation, samples, logistics, metadata curation and software and analysis tools:**

Tanya Golubchik ^27^.

**Leadership, supervision, metadata curation, project administration, samples, logistics sequencing and analysis:**

M. Estee Torok ^8, 10^.

**Project administration, metadata curation, samples, logistics, sequencing, analysis, and software and analysis tools:**

William L Hamilton ^8, 10^.

**Leadership, supervision, samples logistics, project administration, funding acquisition sequencing and analysis:**

David Bonsall ^27^.

**Leadership and supervision, sequencing, analysis, funding acquisition, visualisation and software and analysis tools:**

Ali R Awan ^74^.

**Leadership and supervision, funding acquisition, sequencing, analysis, metadata curation, samples and logistics:**

Sally Corden^33^.

**Leadership supervision, sequencing analysis, samples, logistics, and metadata curation:**

Ian Goodfellow ^11^.

**Leadership, supervision, sequencing, analysis, samples, logistics, and Project administration:**

Darren L Smith ^60, 61^.

**Project administration, metadata curation, samples, logistics, sequencing and analysis:**

Martin D Curran ^14^, and Surendra Parmar ^14^.

**Samples, logistics, metadata curation, project administration sequencing and analysis:**

James G Shepherd ^21^.

**Sequencing, analysis, project administration, metadata curation and software and analysis tools:**

Matthew D Parker ^38^ and Dinesh Aggarwal ^1, 2, 3^.

**Leadership, supervision, funding acquisition, samples, logistics, and metadata curation:**

Catherine Moore ^33^.

**Leadership, supervision, metadata curation, samples, logistics, sequencing and analysis:**

Derek J Fairley^6, 88^, Matthew W Loose ^54^, and Joanne Watkins ^33^.

**Metadata curation, sequencing, analysis, leadership, supervision and software and analysis tools:**

Matthew Bull ^33^, and Sam Nicholls ^15^.

**Leadership, supervision, visualisation, sequencing, analysis and software and analysis tools:**

David M Aanensen ^1, 30^.

**Sequencing, analysis, samples, logistics, metadata curation, and visualisation:**

Sharon Glaysher ^70^.

**Metadata curation, sequencing, analysis, visualisation, software and analysis tools:**

Matthew Bashton ^60^, and Nicole Pacchiarini ^33^.

**Sequencing, analysis, visualisation, metadata curation, and software and analysis tools**: Anthony P Underwood ^1, 30^.

**Funding acquisition, leadership, supervision and project administration:**

Thushan I de Silva ^38^, and Dennis Wang ^38^.

**Project administration, samples, logistics, leadership and supervision**:

Monique Andersson^28^, Anoop J Chauhan ^70^, Mariateresa de Cesare ^26^, Catherine Ludden ^1,3^, and Tabitha W Mahungu ^91^.

**Sequencing, analysis, project administration and metadata curation:**

Rebecca Dewar ^20^, and Martin P McHugh ^20^.

**Samples, logistics, metadata curation and project administration:**

Natasha G Jesudason ^21^, Kathy K Li MBBCh ^21^, Rajiv N Shah ^21^, and Yusri Taha ^66^.

**Leadership, supervision, funding acquisition and metadata curation:**

Kate E Templeton ^20^.

**Leadership, supervision, funding acquisition, sequencing and analysis:**

Simon Cottrell ^33^, Justin O’Grady ^51^, Andrew Rambaut ^19^, and Colin P Smith^93^.

**Leadership, supervision, metadata curation, sequencing and analysis:**

Matthew T.G. Holden ^87^, and Emma C Thomson ^21^.

**Leadership, supervision, samples, logistics and metadata curation**: Samuel Moses ^81, 82^.

**Sequencing, analysis, leadership, supervision, samples and logistics:**

Meera Chand ^7^, Chrystala Constantinidou ^71^, Alistair C Darby ^46^, Julian A Hiscox ^46^, Steve Paterson ^46^, and Meera Unnikrishnan ^71^.

**Sequencing, analysis, leadership and supervision and software and analysis tools:**

Andrew J Page ^51^, and Erik M Volz ^96^.

**Samples, logistics, sequencing, analysis and metadata curation:**

Charlotte J Houldcroft ^8^, Aminu S Jahun ^11^, James P McKenna ^88^, Luke W Meredith ^11^, Andrew Nelson ^61^, Sarojini Pandey ^72^, and Gregory R Young ^60^.

**Sequencing, analysis, metadata curation, and software and analysis tools:**

Anna Price ^34^, Sara Rey ^33^, Sunando Roy ^41^, Ben Temperton^49^, and Matthew Wyles ^38^.

**Sequencing, analysis, metadata curation and visualisation:**

Stefan Rooke^19^, and Sharif Shaaban ^87^.

**Visualisation, sequencing, analysis and software and analysis tools:**

Helen Adams ^35^, Yann Bourgeois ^69^, Katie F Loveson ^68^, Áine O’Toole ^19^, and Richard Stark ^71^.

**Project administration, leadership and supervision:**

Ewan M Harrison ^1, 3^, David Heyburn ^33^, and Sharon J Peacock ^2, 3^

**Project administration and funding acquisition:**

David Buck ^26^, and Michaela John^36^

**Sequencing, analysis and project administration:**

Dorota Jamrozy ^1^, and Joshua Quick ^15^

**Samples, logistics, and project administration:**

Rahul Batra ^78^, Katherine L Bellis ^1, 3^, Beth Blane ^3^, Sophia T Girgis ^3^, Angie Green ^26^, Anita Justice ^28^, Mark Kristiansen ^41^, and Rachel J Williams ^41^.

**Project administration, software and analysis tools:**

Radoslaw Poplawski^15^.

**Project administration and visualisation:**

Garry P Scarlett ^69^.

**Leadership, supervision, and funding acquisition:**

John A Todd ^26^, Christophe Fraser ^27^, Judith Breuer ^40,41^, Sergi Castellano ^41^, Stephen L Michell ^49^, Dimitris Gramatopoulos ^73^, and Jonathan Edgeworth ^78^.

**Leadership, supervision and metadata curation:**

Gemma L Kay ^51^.

**Leadership, supervision, sequencing and analysis:**

Ana da Silva Filipe ^21^, Aaron R Jeffries ^49^, Sascha Ott ^71^, Oliver Pybus ^24^, David L Robertson ^21^, David A Simpson ^6^, and Chris Williams ^33^.

**Samples, logistics, leadership and supervision:**

Cressida Auckland ^50^, John Boyes ^83^, Samir Dervisevic ^52^, Sian Ellard ^49, 50^, Sonia Goncalves^1^, Emma J Meader ^51^, Peter Muir ^2^, Husam Osman ^95^, Reenesh Prakash ^52^, Venkat Sivaprakasam ^18^, and Ian B Vipond ^2^.

**Leadership, supervision and visualisation**

Jane AH Masoli ^49, 50^.

**Sequencing, analysis and metadata curation**

Nabil-Fareed Alikhan ^51^, Matthew Carlile ^54^, Noel Craine ^33^, Sam T Haldenby ^46^, Nadine Holmes ^54^, Ronan A Lyons ^37^, Christopher Moore ^54^, Malorie Perry ^33^, Ben Warne ^80^, and Thomas Williams ^19^.

**Samples, logistics and metadata curation:**

Lisa Berry ^72^, Andrew Bosworth ^95^, Julianne Rose Brown ^40^, Sharon Campbell ^67^, Anna Casey ^17^, Gemma Clark ^56^, Jennifer Collins ^66^, Alison Cox ^43, 44^, Thomas Davis ^84^, Gary Eltringham ^66^, Cariad Evans ^38, 39^, Clive Graham ^64^, Fenella Halstead ^18^, Kathryn Ann Harris ^40^, Christopher Holmes ^58^, Stephanie Hutchings ^2^, Miren Iturriza-Gomara ^46^, Kate Johnson ^38, 39^, Katie Jones ^72^, Alexander J Keeley ^38^, Bridget A Knight ^49, 50^, Cherian Koshy^90^, Steven Liggett ^63^, Hannah Lowe ^81^, Anita O Lucaci ^46^, Jessica Lynch ^25, 29^, Patrick C McClure ^55^, Nathan Moore ^31^, Matilde Mori ^25, 29, 32^, David G Partridge ^38, 39^, Pinglawathee Madona ^43, 44^, Hannah M Pymont ^2^, Paul Anthony Randell ^43, 44^, Mohammad Raza ^38, 39^, Felicity Ryan ^81^, Robert Shaw ^28^, Tim J Sloan ^57^, and Emma Swindells ^65^.

**Sequencing, analysis, Samples and logistics:**

Alexander Adams ^33^, Hibo Asad ^33^, Alec Birchley ^33^, Tony Thomas Brooks ^41^, Giselda Bucca ^93^, Ethan Butcher ^70^, Sarah L Caddy ^13^, Laura G Caller ^2, 3, 12^, Yasmin Chaudhry ^11^, Jason Coombes ^33^, Michelle Cronin ^33^, Patricia L Dyal ^41^, Johnathan M Evans ^33^, Laia Fina ^33^, Bree Gatica-Wilcox ^33^, Iliana Georgana ^11^, Lauren Gilbert ^33^, Lee Graham ^33^, Danielle C Groves ^38^, Grant Hall ^11^, Ember Hilvers ^33^, Myra Hosmillo ^11^, Hannah Jones ^33^, Sophie Jones ^33^, Fahad A Khokhar ^13^, Sara Kumziene-Summerhayes ^33^, George MacIntyre-Cockett ^26^, Rocio T Martinez Nunez ^94^, Caoimhe McKerr ^33^, Claire McMurray ^15^, Richard Myers ^7^, Yasmin Nicole Panchbhaya ^41^, Malte L Pinckert ^11^, Amy Plimmer ^33^, Joanne Stockton ^15^, Sarah Taylor ^33^, Alicia Thornton ^7^, Amy Trebes ^26^, Alexander J Trotter ^51^, Helena Jane Tutill ^41^, Charlotte A Williams ^41^, Anna Yakovleva ^11^ and Wen C Yew ^62^.

**Sequencing, analysis and software and analysis tools:**

Mohammad T Alam ^71^, Laura Baxter ^71^, Olivia Boyd ^96^, Fabricia F. Nascimento ^96^, Timothy M Freeman ^38^, Lily Geidelberg ^96^, Joseph Hughes ^21^, David Jorgensen ^96^, Benjamin B Lindsey ^38^, Richard J Orton ^21^, Manon Ragonnet-Cronin ^96^ Joel Southgate ^33, 34^, and Sreenu Vattipally ^21^.

**Samples, logistics and software and analysis tools:**

Igor Starinskij ^23^.

**Visualisation and software and analysis tools:**

Joshua B Singer ^21^, Khalil Abudahab ^1, 30^, Leonardo de Oliveira Martins ^51^, Thanh Le-Viet ^51^, Mirko Menegazzo ^30^, Ben EW Taylor ^1, 30^, and Corin A Yeats ^30^.

**Project Administration:**

Sophie Palmer ^3^, Carol M Churcher ^3^, Alisha Davies ^33^, Elen De Lacy ^33^, Fatima Downing ^33^, Sue Edwards ^33^, Nikki Smith ^38^, Francesc Coll ^97^, Nazreen F Hadjirin ^3^ and Frances Bolt ^44, 45^.

**Leadership and supervision:**

Alex Alderton^1^, Matt Berriman^1^, Ian G Charles ^51^, Nicholas Cortes ^31^, Tanya Curran ^88^, John Danesh^1^, Sahar Eldirdiri ^84^, Ngozi Elumogo ^52^, Andrew Hattersley ^49, 50^, Alison Holmes ^44, 45^, Robin Howe ^33^, Rachel Jones ^33^, Anita Kenyon ^84^, Robert A Kingsley ^51^, Dominic Kwiatkowski ^1, 9^, Cordelia Langford^1^, Jenifer Mason^48^, Alison E Mather ^51^, Lizzie Meadows ^51^, Sian Morgan ^36^, James Price ^44, 45^, Trevor I Robinson ^48^, Giri Shankar ^33^, John Wain ^51^, and Mark A Webber ^51^.

**Metadata curation:**

Declan T Bradley ^5, 6^, Michael R Chapman ^1, 3, 4^, Derrick Crooke ^28^, David Eyre ^28^, Martyn Guest ^34^, Huw Gulliver ^34^, Sarah Hoosdally ^28^, Christine Kitchen ^34^, Ian Merrick ^34^, Siddharth Mookerjee ^44, 45^, Robert Munn ^34^, Timothy Peto ^28^, Will Potter ^52^, Dheeraj K Sethi ^52^, Wendy Smith ^56^, Luke B Snell ^75, 94^, Rachael Stanley ^52^, Claire Stuart ^52^ and Elizabeth Wastenge^20^.

**Sequencing and analysis:**

Erwan Acheson ^6^, Safiah Afifi ^36^, Elias Allara ^2, 3^, Roberto Amato ^1^, Adrienn Angyal ^38^, Elihu Aranday-Cortes ^21^, Cristina Ariani ^1^, Jordan Ashworth ^19^, Stephen Attwood ^24^, Alp Aydin ^51^, David J Baker ^51^, Carlos E Balcazar ^19^, Angela Beckett ^68^ Robert Beer ^36^, Gilberto Betancor ^76^, Emma Betteridge ^1^, David Bibby ^7^, Daniel Bradshaw^7^, Catherine Bresner ^34^, Hannah E Bridgewater ^71^, Alice Broos ^21^, Rebecca Brown ^38^, Paul E Brown ^71^, Kirstyn Brunker ^22^, Stephen N Carmichael ^21^, Jeffrey K. J. Cheng ^71^, Dr Rachel Colquhoun ^19^, Gavin Dabrera ^7^, Johnny Debebe ^54^, Eleanor Drury ^1^, Louis du Plessis ^24^, Richard Eccles ^46^, Nicholas Ellaby ^7^, Audrey Farbos ^49^, Ben Farr ^1^, Jacqueline Findlay ^41^, Chloe L Fisher ^74^, Leysa Marie Forrest ^41^, Sarah Francois ^24^, Lucy R. Frost ^71^, William Fuller^34^, Eileen Gallagher ^7^, Michael D Gallagher ^19^, Matthew Gemmell ^46^, Rachel AJ Gilroy ^51^, Scott Goodwin ^1^, Luke R Green ^38^, Richard Gregory ^46^, Natalie Groves ^7^, James W Harrison ^49^, Hassan Hartman ^7^, Andrew R Hesketh ^93^, Verity Hill ^19^, Jonathan Hubb ^7^, Margaret Hughes^46^, David K Jackson ^1^, Ben Jackson ^19^, Keith James ^1^, Natasha Johnson ^21^, Ian Johnston ^1^, Jon-Paul Keatley ^1^, Moritz Kraemer ^24^, Angie Lackenby ^7^, Mara Lawniczak ^1^, David Lee ^7^, Rich Livett ^1^, Stephanie Lo ^1^, Daniel Mair ^21^, Joshua Maksimovic ^36^, Nikos Manesis ^7^, Robin Manley ^49^, Carmen Manso ^7^, Angela Marchbank ^34^, Inigo Martincorena ^1^, Tamyo Mbisa ^7^, Kathryn McCluggage ^36^, JT McCrone ^19^, Shahjahan Miah ^7^, Michelle L Michelsen ^49^, Mari Morgan ^33^, Gaia Nebbia ^78^, Charlotte Nelson ^46^, Jenna Nichols ^21^, Paola Niola ^41^, Kyriaki Nomikou ^21^, Steve Palmer ^1^, Naomi Park ^1^, Yasmin A Parr ^1^, Paul J Parsons ^38^, Vineet Patel ^7^, Minal Patel ^1^, Clare Pearson ^2, 1^, Steven Platt ^7^, Christoph Puethe ^1^, Mike Quail ^1^, Jayna Raghwani ^24^, Lucille Rainbow ^46^, Shavanthi Rajatileka ^1^, Mary Ramsay ^7^, Paola C Resende Silva ^41, 42^, Steven Rudder 51, Chris Ruis ^3^, Christine M Sambles ^49^, Fei Sang ^54^, Ulf Schaefer^7^, Emily Scher ^19^, Carol Scott ^1^, Lesley Shirley ^1^, Adrian W Signell ^76^, John Sillitoe ^1^, Christen Smith ^1^, Dr Katherine L Smollett ^21^, Karla Spellman ^36^, Thomas D Stanton ^19^, David J Studholme ^49^, Grace Taylor-Joyce ^71^, Ana P Tedim ^51^, Thomas Thompson ^6^, Nicholas M Thomson ^51^, Scott Thurston^1^, Lily Tong ^21^, Gerry Tonkin-Hill ^1^, Rachel M Tucker ^38^, Edith E Vamos ^4^, Tetyana Vasylyeva^24^, Joanna Warwick-Dugdale ^49^, Danni Weldon ^1^, Mark Whitehead ^46^, David Williams ^7^, Kathleen A Williamson ^19^, Harry D Wilson ^76^, Trudy Workman ^34^, Muhammad Yasir^51^, Xiaoyu Yu ^19^, and Alex Zarebski ^24^.

**Samples and logistics:**

Evelien M Adriaenssens ^51^, Shazaad S Y Ahmad ^2, 47^, Adela Alcolea-Medina ^59, 77^, John Allan ^60^, Patawee Asamaphan ^21^, Laura Atkinson ^40^, Paul Baker ^63^, Jonathan Ball ^55^, Edward Barton^64^, Mathew A Beale^1^, Charlotte Beaver^1^, Andrew Beggs ^16^, Andrew Bell ^51^, Duncan J Berger ^1^, Louise Berry. ^56^, Claire M Bewshea ^49^, Kelly Bicknell ^70^, Paul Bird ^58^, Chloe Bishop ^7^, Tim Boswell ^56^, Cassie Breen ^48^, Sarah K Buddenborg^1^, Shirelle Burton-Fanning ^66^, Vicki Chalker ^7^, Joseph G Chappell ^55^, Themoula Charalampous ^78, 94^, Claire Cormie^3^, Nick Cortes^29, 25^, Lindsay J Coupland ^52^, Angela Cowell ^48^, Rose K Davidson ^53^, Joana Dias ^3^, Maria Diaz ^51^, Thomas Dibling^1^, Matthew J Dorman^1^, Nichola Duckworth^57^, Scott Elliott^70^, Sarah Essex^63^, Karlie Fallon ^58^, Theresa Feltwell ^8^, Vicki M Fleming ^56^, Sally Forrest ^3^, Luke Foulser^1^, Maria V Garcia-Casado^1^, Artemis Gavriil ^41^, Ryan P George ^47^, Laura Gifford ^33^, Harmeet K Gill ^3^, Jane Greenaway ^65^, Luke Griffith^53^, Ana Victoria Gutierrez^51^, Antony D Hale ^85^, Tanzina Haque ^91^, Katherine L Harper ^85^, Ian Harrison ^7^, Judith Heaney ^89^, Thomas Helmer ^58^, Ellen E Higginson^3^, Richard Hopes ^2^, Hannah C Howson-Wells ^56^, Adam D Hunter ^1^, Robert Impey ^70^, Dianne Irish-Tavares ^91^, David A Jackson^1^, Kathryn A Jackson ^46^, Amelia Joseph ^56^, Leanne Kane ^1^, Sally Kay ^1^, Leanne M Kermack ^3^, Manjinder Khakh ^56^, Stephen P Kidd ^29, 25,31^, Anastasia Kolyva ^51^, Jack CD Lee ^40^, Laura Letchford ^1^, Nick Levene ^79^, Lisa J Levett ^89^, Michelle M Lister ^56^, Allyson Lloyd ^70^, Joshua Loh ^60^, Louissa R Macfarlane-Smith ^85^, Nicholas W Machin ^2, 47^, Mailis Maes ^3^, Samantha McGuigan ^1^, Liz McMinn ^1^, Lamia Mestek-Boukhibar ^41^, Zoltan Molnar ^6^, Lynn Monaghan ^79^, Catrin Moore ^27^, Plamena Naydenova ^3^, Alexandra S Neaverson ^1^, Rachel Nelson ^1^, Marc O Niebel ^21^, Elaine O’Toole^48^, Debra Padgett ^64^, Gaurang Patel ^1^, Brendan AI Payne ^66^, Liam Prestwood ^1^, Veena Raviprakash ^67^, Nicola Reynolds^86^, Alex Richter ^16^, Esther Robinson ^95^, Hazel A Rogers^1^, Aileen Rowan ^96^, Garren Scott ^64^, Divya Shah ^40^, Nicola Sheriff ^67^, Graciela Sluga, Emily Souster^1^, Michael Spencer-Chapman^1^, Sushmita Sridhar ^1, 3^, Tracey Swingler ^53^, Julian Tang^58^, Graham P Taylor^96^, Theocharis Tsoleridis ^55^, Lance Turtle^46^, Sarah Walsh ^57^, Michelle Wantoch ^86^, Joanne Watts ^48^, Sheila Waugh ^66^, Sam Weeks^41^, Rebecca Williams^31^, Iona Willingham^56^, Emma L Wise ^25, 29, 31^, Victoria Wright ^54^, Sarah Wyllie ^70^, and Jamie Young ^3^.

**Software and analysis tools**

Amy Gaskin^33^, Will Rowe ^15^, and Igor Siveroni ^96^.

**Visualisation**

Robert Johnson ^96^.

**1** Wellcome Sanger Institute, **2** Public Health England, **3** University of Cambridge, **4** Health Data Research UK, Cambridge, **5** Public Health Agency, Northern Ireland, **6** Queen’s University Belfast **7** Public Health England Colindale, **8** Department of Medicine, University of Cambridge, **9** University of Oxford, **10** Departments of Infectious Diseases and Microbiology, Cambridge University Hospitals NHS Foundation Trust; Cambridge, UK, **11** Division of Virology, Department of Pathology, University of Cambridge, **12** The Francis Crick Institute, **13** Cambridge Institute for Therapeutic Immunology and Infectious Disease, Department of Medicine, **14** Public Health England, Clinical Microbiology and Public Health Laboratory, Cambridge, UK, **15** Institute of Microbiology and Infection, University of Birmingham, **16** University of Birmingham, **17** Queen Elizabeth Hospital, **18** Heartlands Hospital, **19** University of Edinburgh, **20** NHS Lothian, **21** MRC-University of Glasgow Centre for Virus Research, **22** Institute of Biodiversity, Animal Health & Comparative Medicine, University of Glasgow, **23** West of Scotland Specialist Virology Centre, **24** Dept Zoology, University of Oxford, **25** University of Surrey, **26** Wellcome Centre for Human Genetics, Nuffield Department of Medicine, University of Oxford, **27** Big Data Institute, Nuffield Department of Medicine, University of Oxford, **28** Oxford University Hospitals NHS Foundation Trust, **29** Basingstoke Hospital, **30** Centre for Genomic Pathogen Surveillance, University of Oxford, **31** Hampshire Hospitals NHS Foundation Trust, **32** University of Southampton, **33** Public Health Wales NHS Trust, **34** Cardiff University, **35** Betsi Cadwaladr University Health Board, **36** Cardiff and Vale University Health Board, **37** Swansea University, **38** University of Sheffield, **39** Sheffield Teaching Hospitals, **40** Great Ormond Street NHS Foundation Trust, **41** University College London, **42** Oswaldo Cruz Institute, Rio de Janeiro **43** North West London Pathology, **44** Imperial College Healthcare NHS Trust, **45** NIHR Health Protection Research Unit in HCAI and AMR, Imperial College London, **46** University of Liverpool, **47** Manchester University NHS Foundation Trust, **48** Liverpool Clinical Laboratories, **49** University of Exeter, **50** Royal Devon and Exeter NHS Foundation Trust, **51** Quadram Institute Bioscience, University of East Anglia, **52** Norfolk and Norwich University Hospital, **53** University of East Anglia, **54** Deep Seq, School of Life Sciences, Queens Medical Centre, University of Nottingham, **55** Virology, School of Life Sciences, Queens Medical Centre, University of Nottingham, **56** Clinical Microbiology Department, Queens Medical Centre, **57** PathLinks, Northern Lincolnshire & Goole NHS Foundation Trust, **58** Clinical Microbiology, University Hospitals of Leicester NHS Trust, **59** Viapath, **60** Hub for Biotechnology in the Built Environment, Northumbria University, **61** NU-OMICS Northumbria University, **62** Northumbria University, **63** South Tees Hospitals NHS Foundation Trust, **64** North Cumbria Integrated Care NHS Foundation Trust, **65** North Tees and Hartlepool NHS Foundation Trust, **66** Newcastle Hospitals NHS Foundation Trust, **67** County Durham and Darlington NHS Foundation Trust, **68** Centre for Enzyme Innovation, University of Portsmouth, **69** School of Biological Sciences, University of Portsmouth, **70** Portsmouth Hospitals NHS Trust, **71** University of Warwick, **72** University Hospitals Coventry and Warwickshire, **73** Warwick Medical School and Institute of Precision Diagnostics, Pathology, UHCW NHS Trust, **74** Genomics Innovation Unit, Guy’s and St. Thomas’ NHS Foundation Trust, **75** Centre for Clinical Infection & Diagnostics Research, St. Thomas’ Hospital and Kings College London, **76** Department of Infectious Diseases, King’s College London, **77** Guy’s and St. Thomas’ Hospitals NHS Foundation Trust, **78** Centre for Clinical Infection and Diagnostics Research, Department of Infectious Diseases, Guy’s and St Thomas’ NHS Foundation Trust, **79** Princess Alexandra Hospital Microbiology Dept., **80** Cambridge University Hospitals NHS Foundation Trust, **81** East Kent Hospitals University NHS Foundation Trust, **82** University of Kent, **83** Gloucestershire Hospitals NHS Foundation Trust, **84** Department of Microbiology, Kettering General Hospital, **85** National Infection Service, PHE and Leeds Teaching Hospitals Trust, **86** Cambridge Stem Cell Institute, University of Cambridge, **87** Public Health Scotland, 88 Belfast Health & Social Care Trust, **89** Health Services Laboratories, **90** Barking, Havering and Redbridge University Hospitals NHS Trust, **91** Royal Free NHS Trust, **92** Maidstone and Tunbridge Wells NHS Trust, **93** University of Brighton, **94** Kings College London, **95** PHE Heartlands, **96** Imperial College London, **97** Department of Infection Biology, London School of Hygiene and Tropical Medicine.

## Supplementary Information

**Figure S1.**
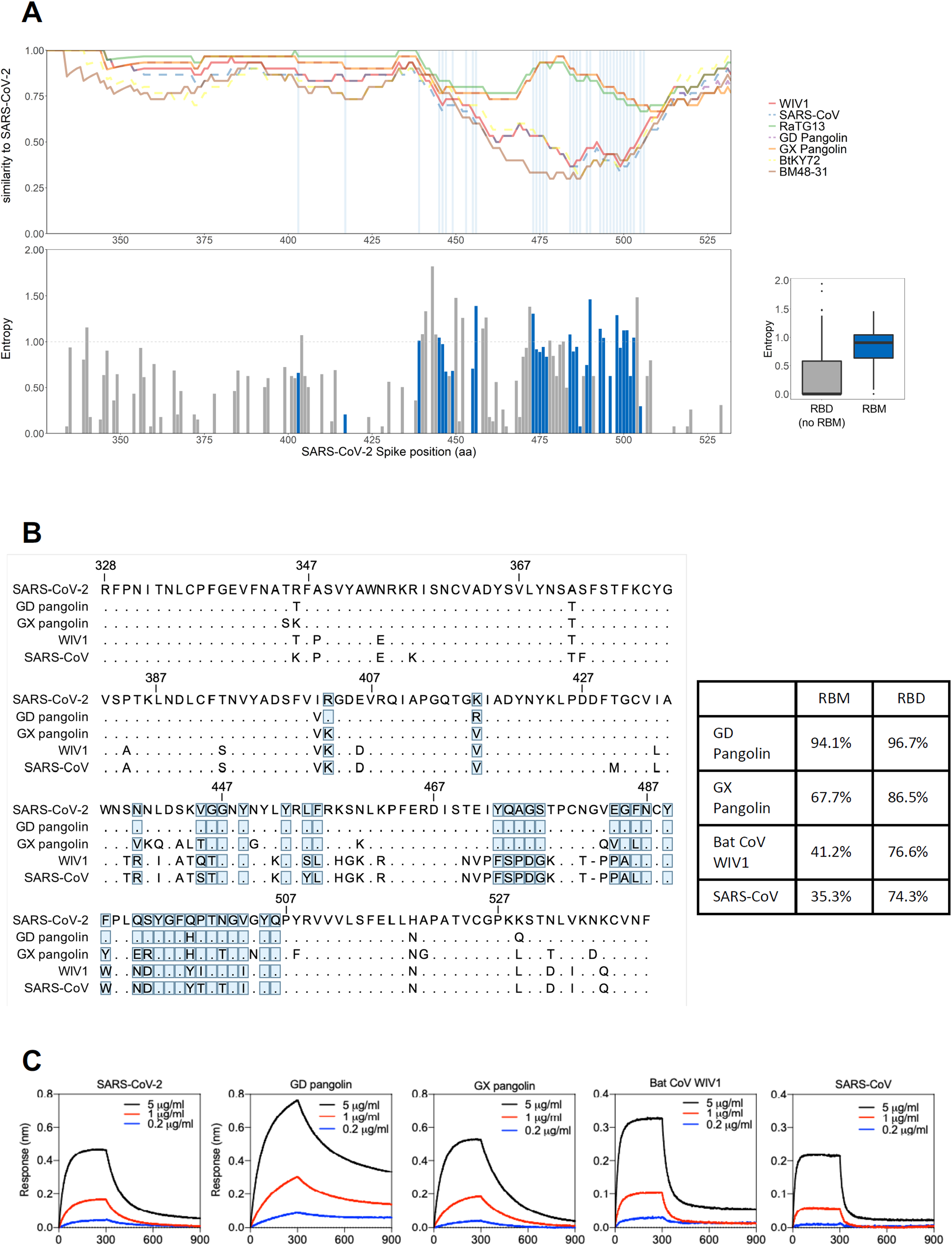
RBDs from bat and pangolin *Sarbecovirus* isolates bind to hACE2 despite RBM divergence. **(A)** Top – Pairwise similarity to SARS-CoV-2 (sliding window size of 30 amino acids) for seven related *Sarbecoviruses* (see figure key) across the RBD region of the Spike protein. Bottom – Site-specific entropy plot across the RBD protein alignment of SARS-CoV-2 and 68 related viruses (**Data S1**). Entropy for each position *l* (H(*l*)) was calculated using Shannon’s entropy formula with a natural log as implemented in Bioedit (H(*l*) = -Σf(*a,l*)ln(f(*a,l*)); f(*a,l*) being the frequency of amino acid *a* at position *l*). Sites constituting the RBM are annotated in blue; the x-axis refers to absolute positions in the SARS-CoV-2 Spike protein sequence. Right – box plot of site-specific entropy values for the RBM sites (blue) and remaining non-RBM RBD sites (gray). **(B)** Sequence alignment (left) and identity for RBM and RBD (right) to SARS-CoV-2 of the RBD sequences showing binding to hACE2. RBM residues indicated by blue boxes. **(C)** Binding of hACE2 to human, pangolin and bat *Sarbecovirus* RBDs by BLI. Bat CoV RaTG13, Bat CoVs ZC45, BtKY72 and BGR2008 have also been tested and did not bind hACE2.

**Figure S2.**
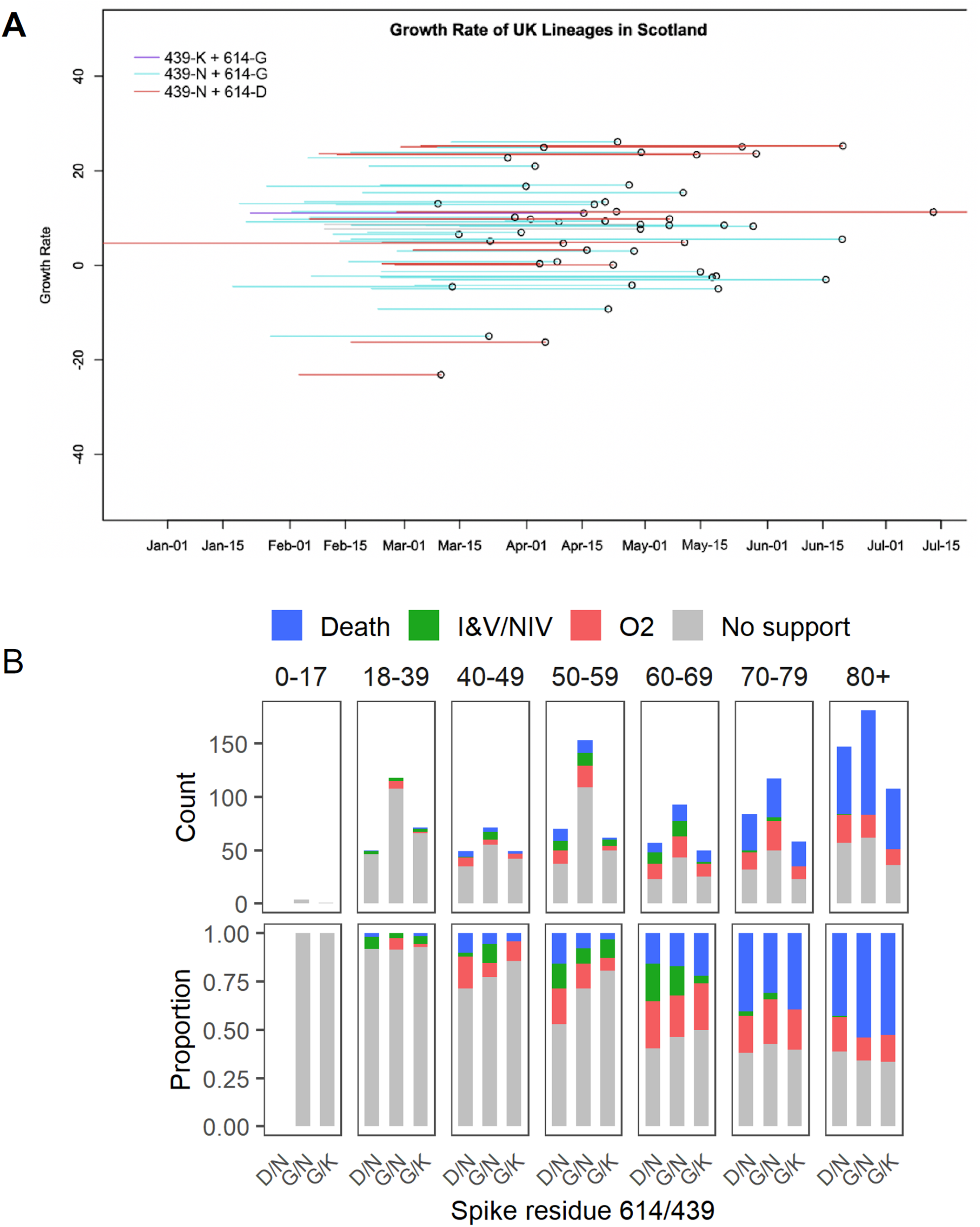
Virological and clinical results stratified by positions 439 and 614. **(A)** Phylodynamic analysis showing lineage growth rates relative to sampling times for UK lineages in Scotland. The Scottish N439K lineage i (which co-occurs with D614G) is indicated in purple along with whether N439 lineages are D614 (red) or D614G (blue). See also “Lineage growth rates” at http://sars2.cvr.gla.ac.uk/RiseFallScotCOVID/. **(B)** Comparison of clinical severity between D614/N439, D614G/N439 and D614G/N439K genotypes by patient age group for 1591 patients whose diagnostic samples were sequenced. Ordinal scale scored by oxygen requirement: 1. No respiratory support, 2: Supplemental oxygen, 3: Invasive or non-invasive ventilation or oxygen delivery by high flow nasal cannulae, 4: Death.

**Figure S3.**
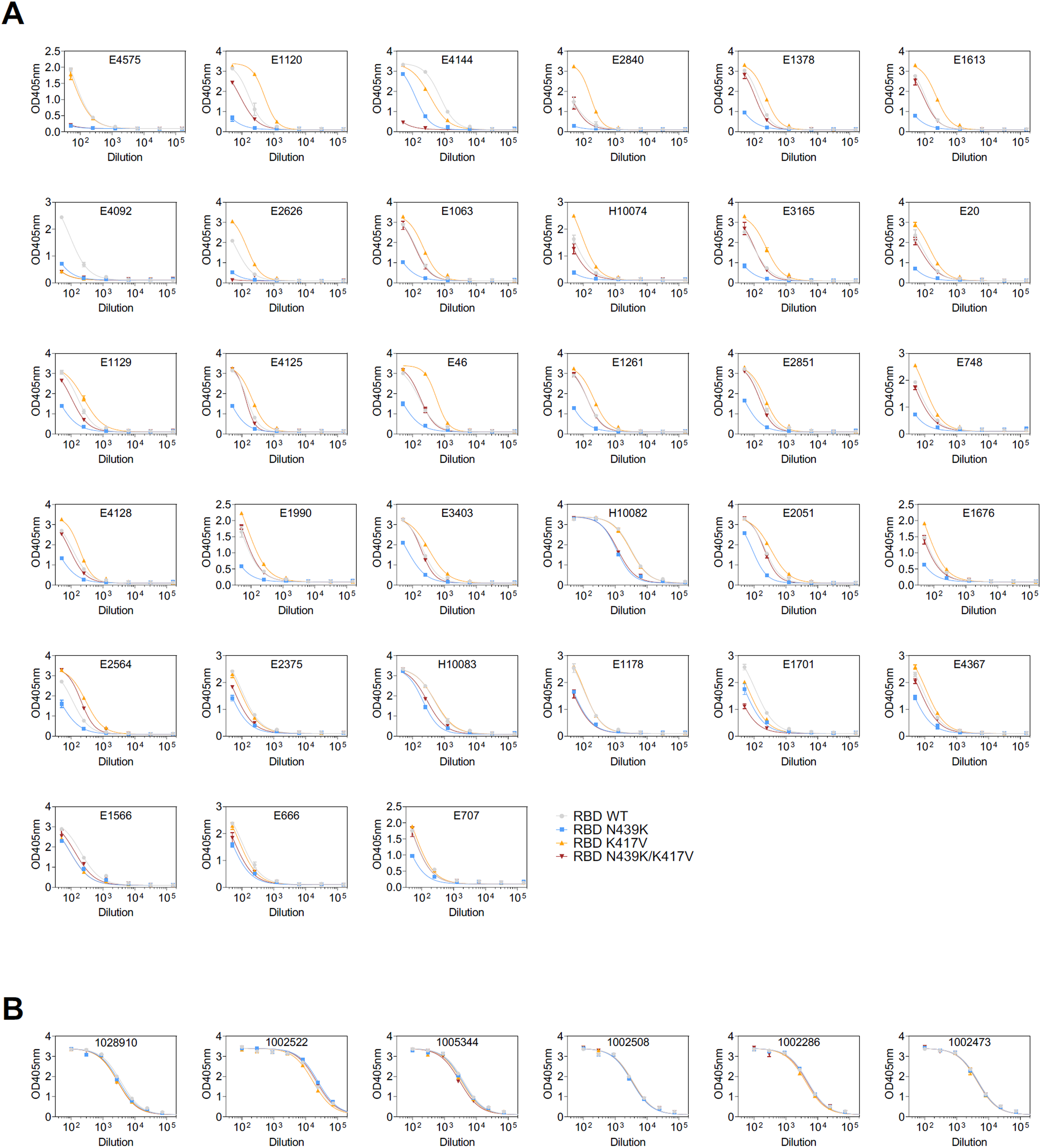
ELISA binding of the 33 human sera with a >2-fold reduction of binding to RBD N439K (A) and of the 6 sera of individuals infected with SARS-CoV-2 N439K variant (B) to RBD WT (grey), N439K (blue), K417V (yellow) and N439K/K417V (red).

**Figure S4.**
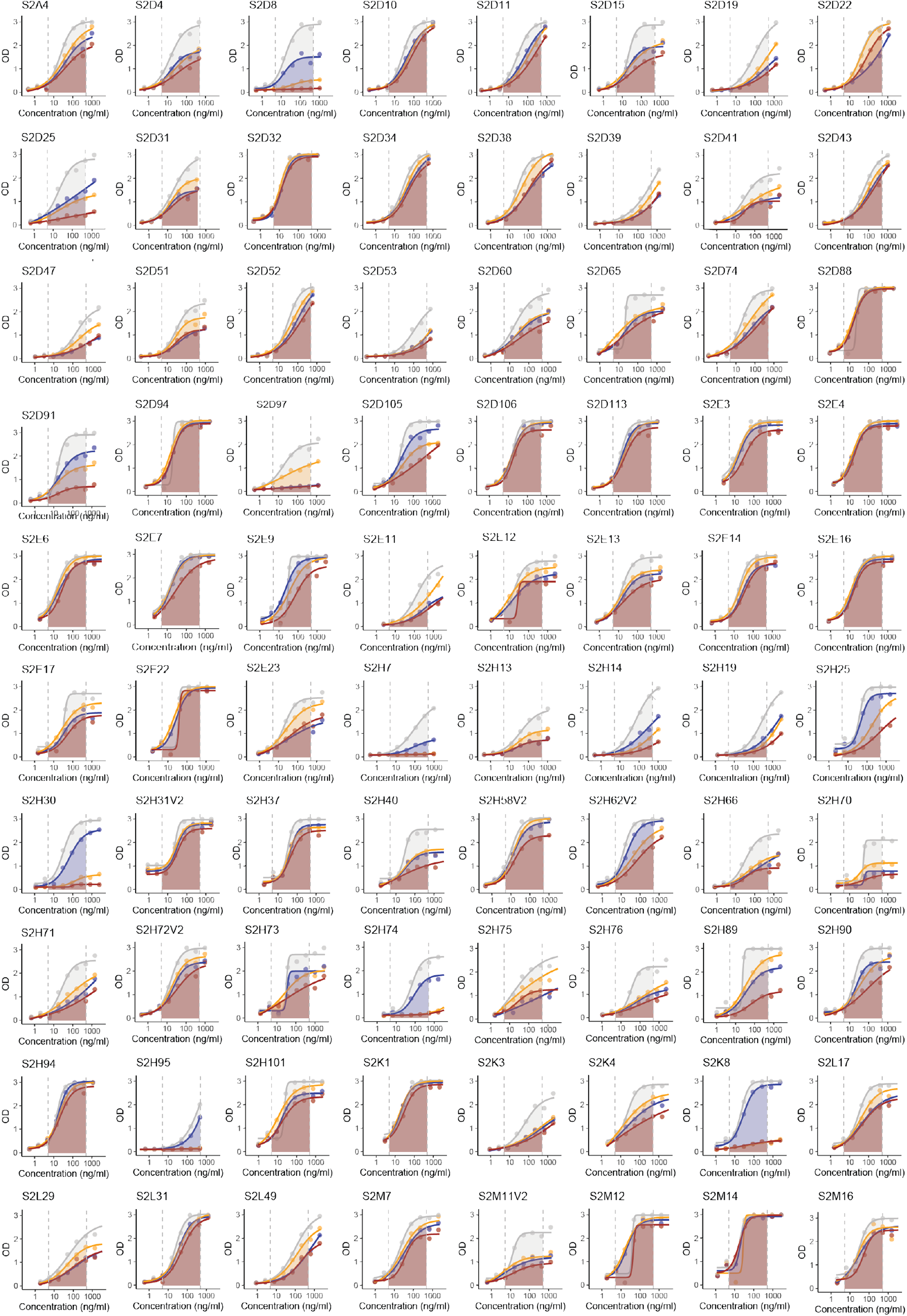
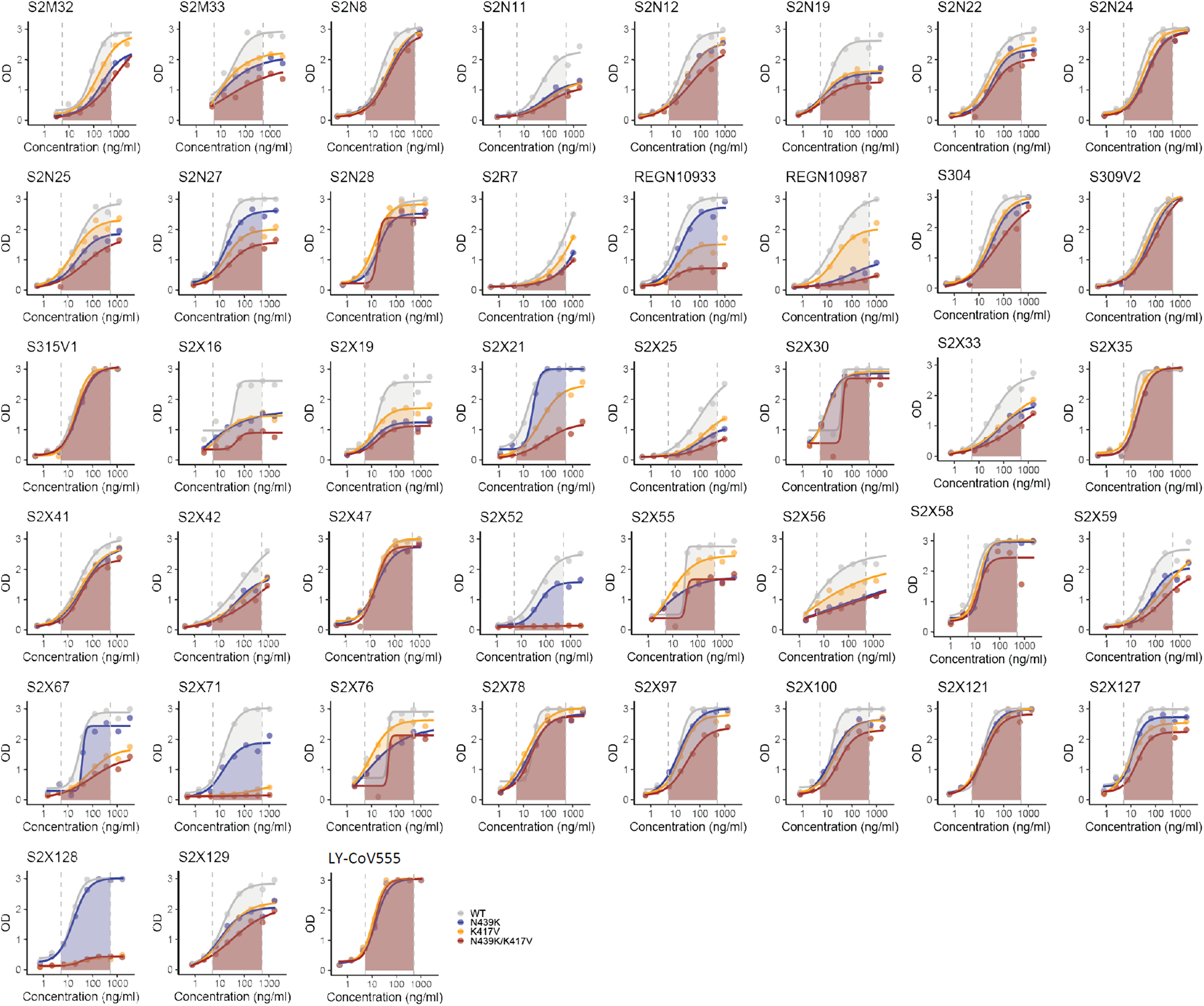
ELISA binding of the 147 mAbs to RBD WT (grey), N439K (blue), K417V (yellow) and N439K/K417V (red). AUC used for quantification is highlighted.

**Figure S5A.**
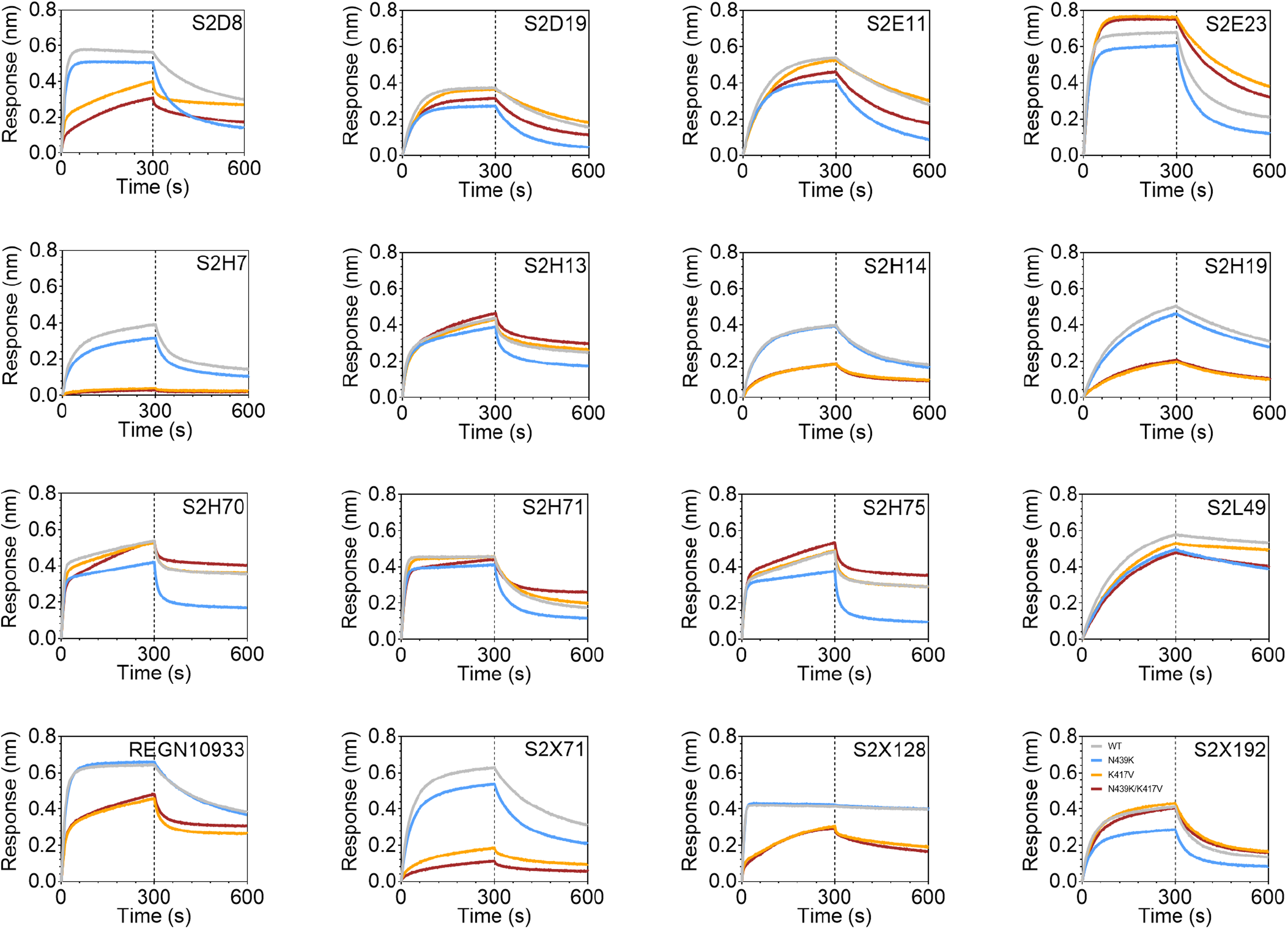
Binding of 15 selected mAbs to RBD WT (grey), N439K (blue), K417V (yellow) and N439K/K417V (red) as measured by BLI.

**Figure S5B.**
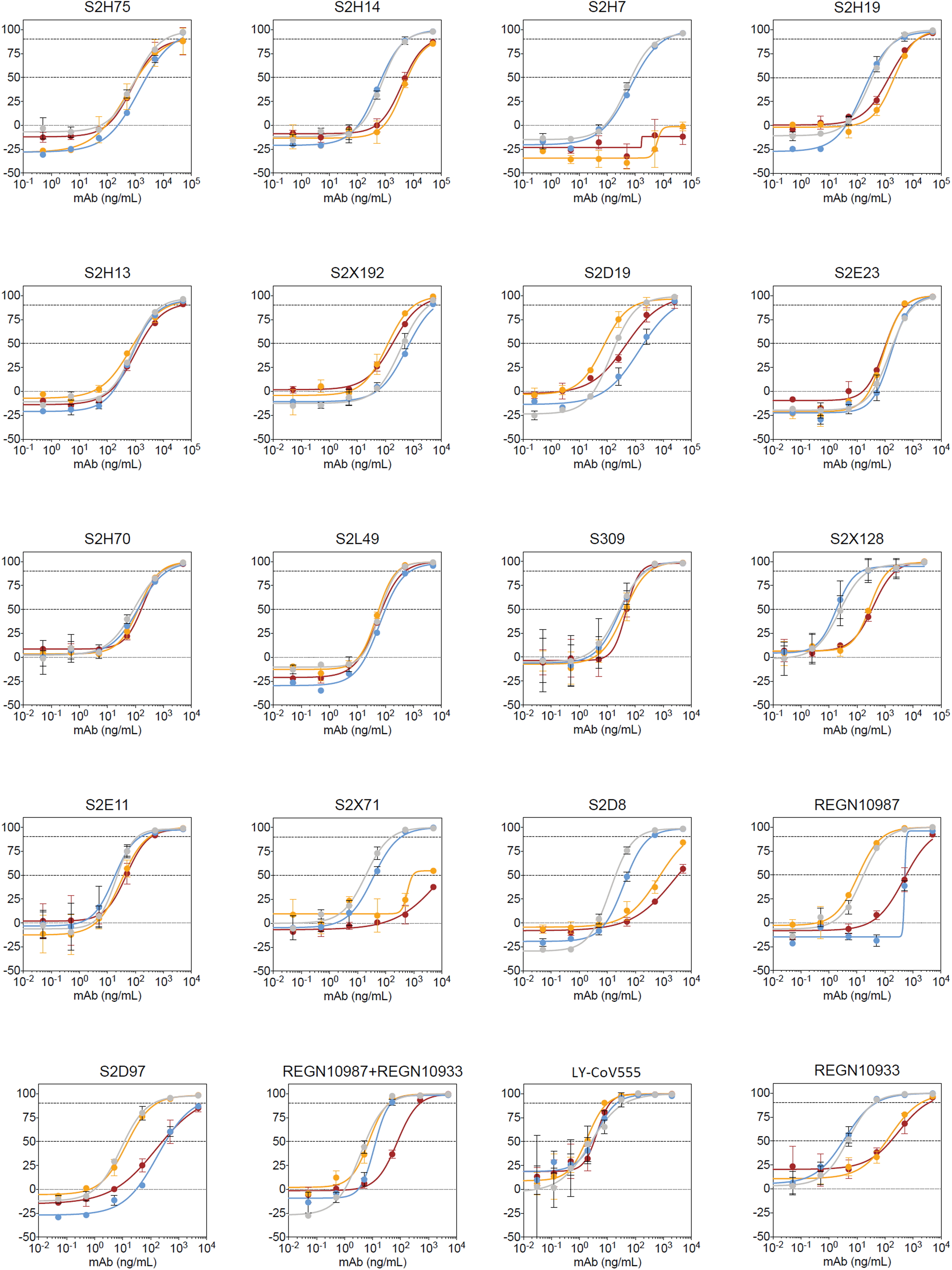
VSV pseudovirus neutralization curves of all mAbs tested. Representative of n=3, bars = STD

**Table S1.**
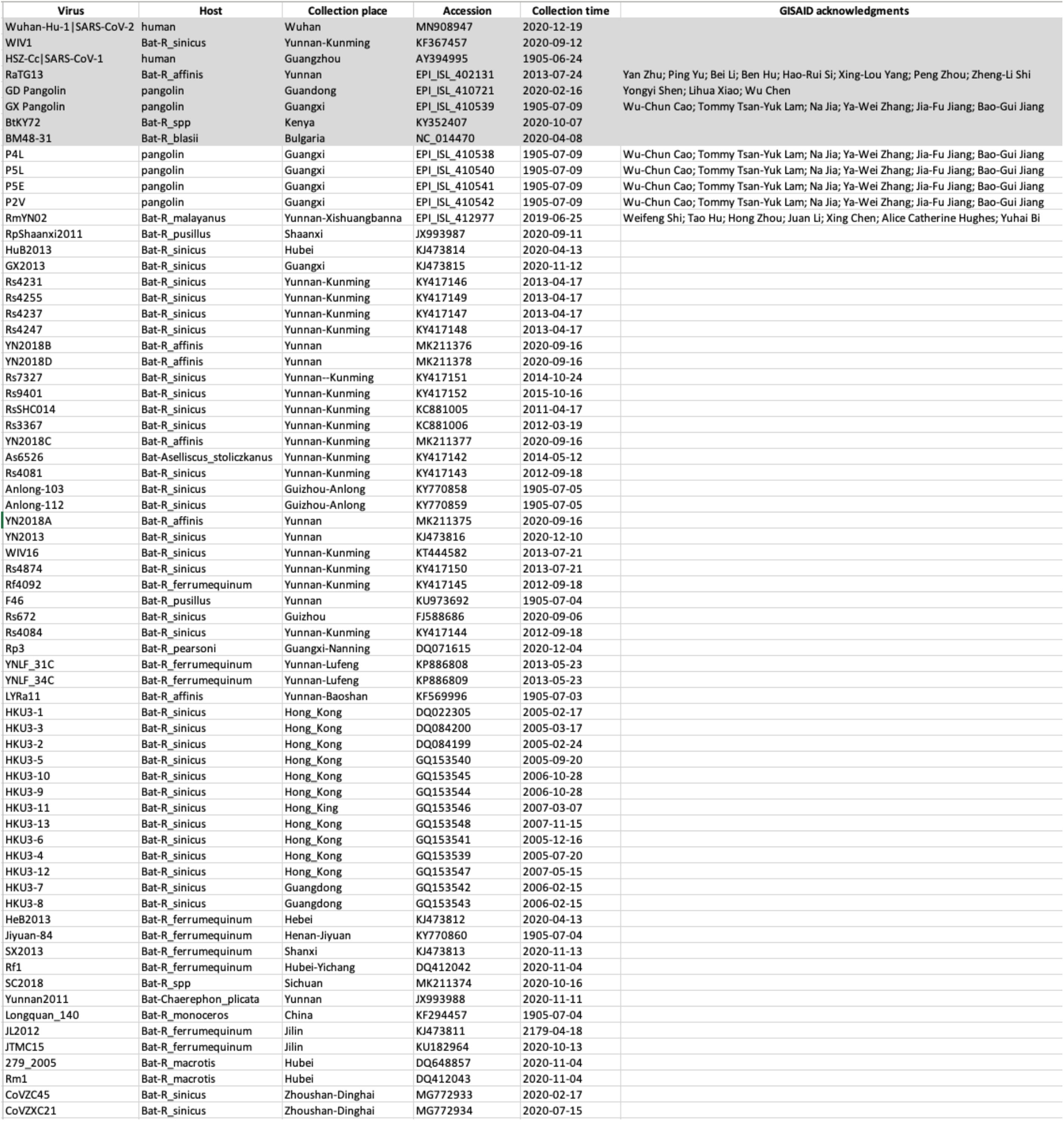
Details of the sarbecovirus sequences used for Figure S1. The top 8 sequences shaded in gray were used for the similarity plot and all 69 sequences were used for the entropy plot.

**Table S2.** Parameter estimates on the link scale from the model estimating the impact of the N439K mutation on the Ct value of patients infected with SARS-CoV-2 in Scotland. Credible intervals represent 95% the shortest posterior density intervals. The difference between D614G/N349 and D614G/N349K was estimated by direct subtraction of the Hamiltonian Monte Carlo samples of the D614G/N349K estimate from the D614G/N349 estimate. Ct value did not appear strongly correlated with biological sex or age after controlling for the other factors. Patients infected with related viral genomes had correlated Ct values at testing potentially implying that there are other undescribed mutations in the genome that are affecting the viral load.

|  | Posterior mean | Lower bound of 95% shortest probability interval | Upper bound of 95% shortest probability interval |
| --- | --- | --- | --- |
| D614G/N349 | -0.31 | -1.91 | 1.48 |
| D614G/N439K | -0.96 | -2.68 | 0.74 |
| Female biological sex | 0.05 | -0.34 | 0.43 |
| Intercept | 27.22 | 26.48 | 27.97 |
| Cluster random effect standard deviation | 0.92 | 0.00 | 1.49 |
| Phylogenetic random effect standard deviation | 0.99 | 0.30 | 1.72 |
| Age penalised spline standard deviation | 1.15 | 0.00 | 3.15 |
| Days since first case in the dataset penalised spline standard deviation | 9.78 | 4.42 | 16.22 |
| Residual standard deviation | 3.91 | 3.71 | 4.12 |
| Difference between D614G/N349 and D614G/N349K | -0.65 | -1.22 | -0.07 |

**Table S3.**
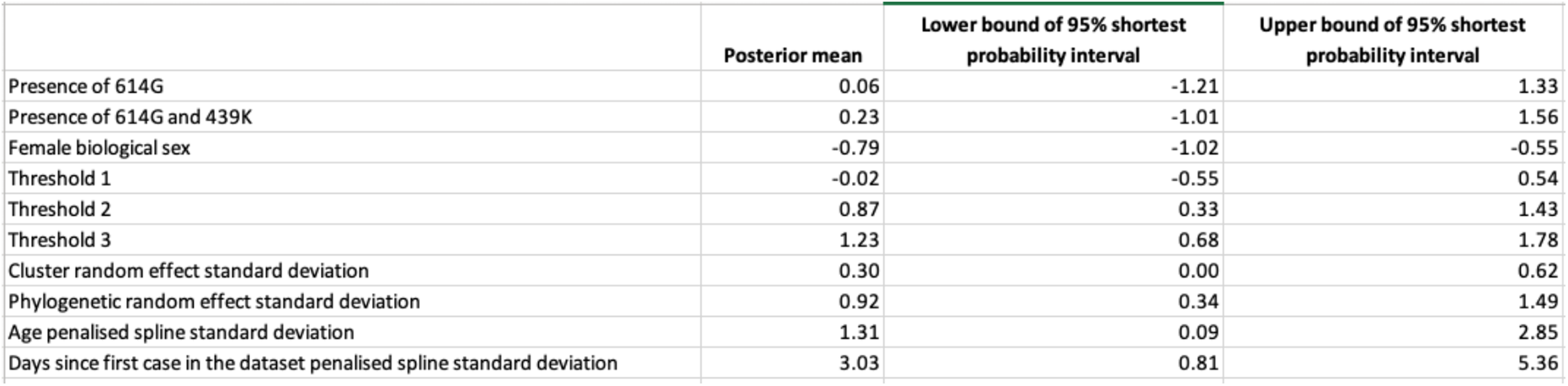
Parameter estimates on the link scale from the model estimating the impact of the N439K mutation on the severity of infection of patients infected with SARS-CoV- 2 in Scotland. Credible intervals represent 95% the shortest posterior density intervals. Thresholds correspond to the positions of the boundaries between the different severity classes.

**Table S4.** Nucleotide Differences between GLA1 and GLA2. SNPs determined by Cov-GLUE on consensus sequences relative to Wuhan-Hu-1 (NC_045512.2).

| Sample | SNP | Amino Acid Change |  |
| --- | --- | --- | --- |
|  |  | Gene | Mutation |
| GLA1 | C3037T | nsp12 | P323L |
|  | C14408T | S | D614G |
|  | A23403G | E | V5A |
|  | A24388T |  |  |
|  | T26258C |  |  |
| GLA2 | C3037T | nsp12 | P323L |
|  | C14408T | nsp15 | V35A |
|  | T19724C | S | N439K |
|  | C22879A | S | D614G |
|  | A23403G | ORF 10 | V6F |
|  | G29573T |  |  |

